# Hi-C profiling in tissues reveals 3D chromatin-regulated breast tumor heterogeneity and tumor-specific looping-mediated biological pathways

**DOI:** 10.1101/2024.03.13.584872

**Authors:** Lavanya Choppavarapu, Kun Fang, Tianxiang Liu, Victor X. Jin

## Abstract

Current knowledge in three-dimensional (3D) chromatin regulation in normal and disease states was mostly accumulated through Hi-C profiling in *in vitro* cell culture system. The limitations include failing to recapitulate disease-specific physiological properties and often lacking clinically relevant disease microenvironment. In this study, we conduct tissue-specific Hi-C profiling in a pilot cohort of 12 breast tissues comprising of two normal tissues (NTs) and ten ER+ breast tumor tissues (TTs) including five primary tumors (PTs), and five tamoxifen-treated recurrent tumors (RTs). We find largely preserved compartments, highly heterogeneous topological associated domains (TADs) and intensively variable chromatin loops among breast tumors, demonstrating 3D chromatin-regulated breast tumor heterogeneity. Further cross-examination identifies RT-specific looping-mediated biological pathways and suggests CA2, an enhancer-promoter looping (EPL)-mediated target gene within the bicarbonate transport metabolism pathway, might play a role in driving the tamoxifen resistance. Remarkably, the inhibition of CA2 not only impedes tumor growth both *in vitro* and *in vivo*, but also reverses chromatin looping. Our study thus yields significant mechanistic insights into the role and clinical relevance of 3D chromatin architecture in breast cancer endocrine resistance.

## INTRODUCTION

Endocrine therapy, including tamoxifen (Tam), is currently a standard and effective treatment for estrogen receptor α positive (ER+) breast cancer patients^**1**^. However, about one third of all ER+ breast cancer patients eventually develop treatment resistance to this hormonal therapy and many even die from this cancer^**2**,**3**^. Current treatment strategy is mainly guided by breast cancer patient prognosis with a typical pathological risk assessment^**4**,**5**^. Such treatment selection generally doesn’t consider individual patient tumor genomic heterogeneity. Recent studies on characterizing the genomic landscape of breast patient cohorts have revealed the molecular differences among individual breast tumors including genomic drivers of breast cancer progression, intra- or inter-tumoral heterogeneity, as well as the variable responses of individual patients to therapies associated with these molecular differences^**6-10**^. Despite these genomic studies of patient cohorts have provided insights into the biological underpinnings of this cancer, the limitation was that they were mainly focused on copy-number and gene expression analyses. Therefore, it is imperative to examine intra- or inter-tumoral heterogeneity in a three-dimensional (3D) genome scale and to determine the difference of 3D chromatin architecture in breast cancer patient cohorts.

Current knowledge in 3D chromatin regulation in normal and disease states was mostly accumulated through Hi-C profiling in *in vitro* cell culture system^**11-16**^. Although these cell model studies have been very useful in unraveling the alteration of 3D-regulated biological pathways in diseases, such studies as many other cell line studies clearly showed many limitations including failing to recapitulate disease-specific physiological properties and often lacking of clinically relevant disease microenvironment. A few recent studies not only adapted Hi-C technique into human patient samples but also revealed patient- or tumor-subtype-specific 3D genome architecture^**17-20**^. For example, Xu et al. identified AML subtype-specific changes in 3D genome features such as compartments, TAD boundaries and chromatin loops and further showed that repressive loops were widespread in the genome^**17**^. Johnstone et al. identified an intermediate compartment between the open A and the closed B compartments that shifted its interactions toward the B compartment in colorectal tumors^**18**^. Kloetgen et al. highlighted 3D landscape variations among different leukemia subtypes and suggested that drugs with reported antileukemic activity partially reversed 3D interactions in specific loci^**19**^. It is becoming very clear that to expand the characterization of 3D genome organization in patient samples is an emerging task. This thus motivates us to conduct this study to elucidate 3D chromatin-regulated breast cancer heterogeneity.

In this study, we conduct a tissue-specific Hi-C profiling^**21**^ in a pilot cohort of 12 breast tissues comprising of two normal tissues (NTs) and ten ER+ breast tumors (TTs) including five primary tumors (PTs), and five tamoxifen-treated recurrent tumors (RTs). Here, we identify differential chromatin compartments among NTs, PTs, and RTs, as well as PT-, RT-, individual tumor (IT)-specific topological associated domains (TADs) and looping genes. We also cross-examine the interplay among the three layers of 3D chromatin structure. We further overlay tumor-derived looping genes and cell-derived looping genes to obtain common looping genes, and then perform *in silico* analyses to identify novel looping-mediated biological pathways that might mediate resistance to endocrine therapy. We finally test an efficacy of an inhibitor targeting metabolism pathways in *in vitro* cell lines and *in vivo* cell-line derived Xenografts (CDXs) and conduct 3C/RT-qPCR validations. Therefore, our study yields significant mechanistic insights into the role and clinical relevance of 3D chromatin architecture in breast cancer endocrine resistance.

## RESULTS

### Compartments are largely preserved among breast tumors

To comprehensively examine the variability of 3D chromatin architecture in breast tumor tissues, we conducted tissue-specific Hi-C profiling in a pilot cohort comprising two NTs, five PTs, and five RTs (**Table S1**) and produced a total of four billion of raw pair-end reads with an average of

∼350 million per tissue (**Figure S1**). We then performed computational analyses to interrogate the interplay of three layers of chromatin architecture in a tumor-specific manner (**Figure 1A**). We first applied HiC-Pro^**22**^ to identify compartment A/B for each of individual tissues and found that there were roughly the same number (∼2,000) of compartments for all 12 tissues and a nearly even number of compartments A and B within each of 12 tissues where a majority of the size of compartments were less than 2 Mb (**Figure 1B**). We then used dcHiC^**23**^ to identify differential compartments between TTs and NTs (TTs *vs* NTs) and between RTs and PTs (RTs *vs* PTs), respectively (**Table S2**). Interestingly, we found 2,085 and 391 bins of 100□Kb/bin were differential compartments at FDR□<□0.1 for TTs *vs* NTs and RTs *vs* PTs, respectively (**Figure 1C**), indicating that the compartment patterns were largely conserved among tumor tissues but exhibited a greater disparity between tumor and normal tissues. We further classified those differential compartment bins into flipping transitions, *i*.*e*., A to B or B to A, and matching transitions, *i*.*e*., A to A or B to B. Of all differential compartment bins between TTs *vs* NTs, 79.6% (46.8% NT A to TT B and 32.8% NT B to TT A) were flipping transitions and 20.4% were matching transitions (**Figure 1D** - top panel and **Figure S2A**). Interestingly, we found that the B-A flipping transition was more dominant in PT to RT transition such that 73.7% (15.3% PT A to RT B and 58.3% PT B to RT A) were flipping transitions and 26.3% were matching transitions (**Figure 1D** - bottom panel and **Figure S2B**). An example of identified flipping and matching transitions between TTs *vs* NTs and RTs *vs* PTs was visualized by IGV (**Figure 1E** and **Figure S2C, D**). Finally, we leveraged a quantile normalized principal component analysis (**METHODS**) to calculate compartment similarities within (intra-) or among (inter-) tissues. We found that the intra-tissue correlations were generally larger than the inter-tissue correlations and the RT-PT correlation was higher than the PT-NT and RT-NT correlations (**Figure 1F**). Taken together, our results demonstrated that 3D chromatin architecture at the compartment layer was largely preserved among breast tumor tissues with a small percentage of chromatin regions undergoing compartment flipping during the progression of breast cancer endocrine resistance.

**Figure 1.**
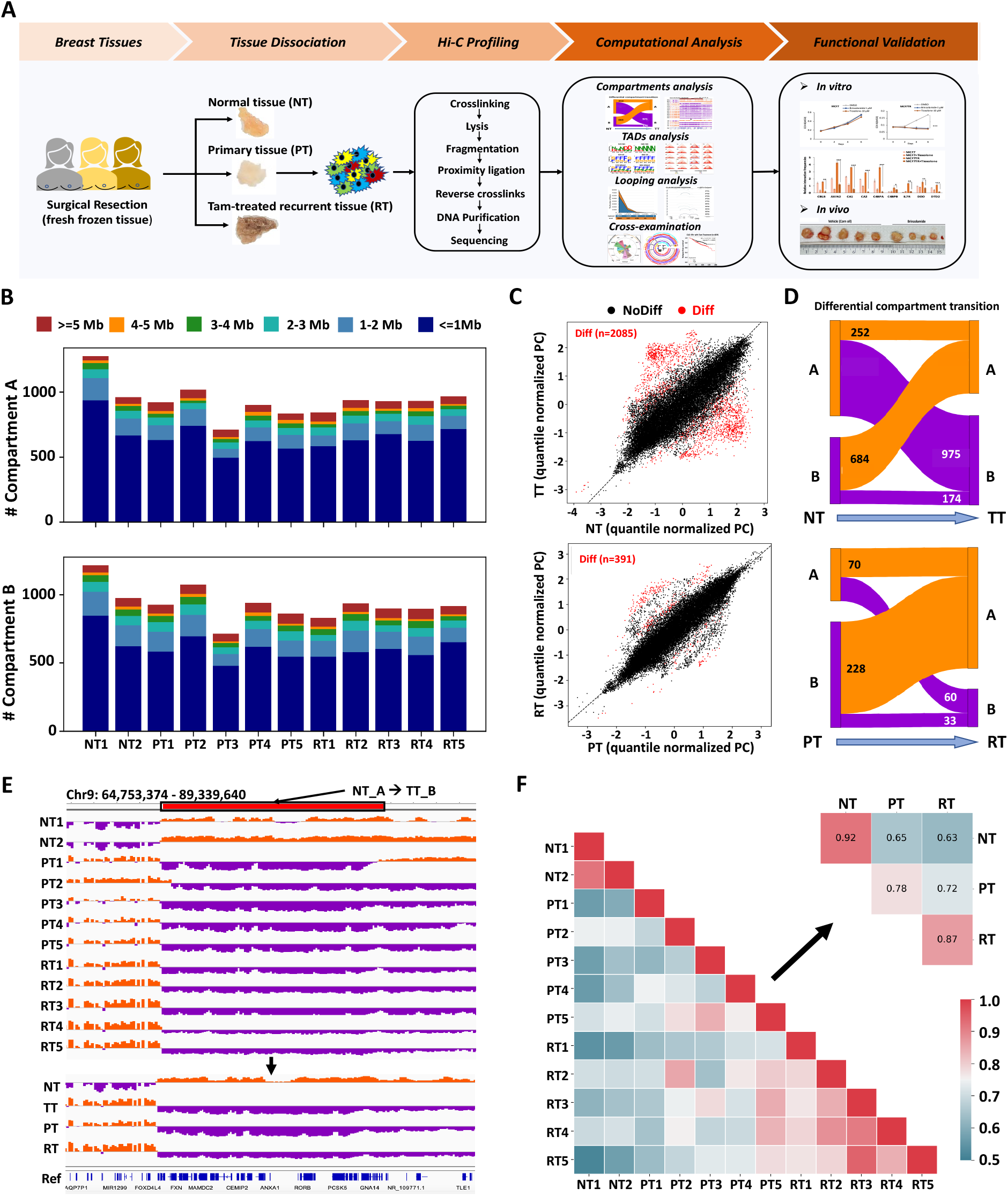
Compartments are largely preserved among breast tumors. **A**. A workflow of the study. Twelve breast normal and tumor tissues were collected, including two NTs, five PTs, and five RTs. Hi-C profiling was then performed for each tissue and generated ∼3 billion reads. Computational analyses were further conducted to investigate the three layers of genome organization, including compartment, TADs and chromatin loops as well as in silico analyses were performed to identify enriched biological pathways and to predict clinical outcome. In vitro and in vivo functional characterization was performed to elicit the looping-mediated biological pathway. **B**. The stacked bar plots showing the number of compartments A and B in different lengths in each of 12 normal and tumor tissues. The number of compartments was around 2,000 across tissues, and the length of the majority of compartment A and B were less than 1Mb. **C**. The scatter plots displaying the number of 100kb bins with statistically differential compartmentalization identified by dcHiC. The number of differential compartments between TT and NT was 2,085, and the number between PT and RT was 391. NT: NT1-2, PT: PT1-5, RT: RT1-5. **D**. The Sankey plots illustrating the breakdown of the numbers of differential compartment calls (100kb resolution) belonging to different types, including flipping (A to B, B to A), and matching (A to A and B to B). Compartment A is in dark-orange, and compartment B is in dark-violet. **E**. An IGV screenshot showing compartment A in NTs transition to compartment B in TTs. **F**. The lower triangle heatmap demonstrating the Pearson correlation of compartmentalization between each tissue sample, and the upper triangle heatmap shows the group-level Pearson correlation within (intra)/ between(inter) NT, PT and RT groups. Generally, intra-group correlation is higher than inter-group correlation. For inter-group correlation, RT-PT correlation is higher than PT-NT and RT-NT groups.

### TADs display a highly heterogeneous among breast tumors

Next, we wanted to examine the tumor-specific chromatin architecture at the TAD layer. We used TopDom^**24**^ to identify ∼6,000 TADs for each of 12 tissues (**Figure 2A** - left panel and **Figure S3A**) and observed a similar size distribution of TADs across the tissues (**Figure 2A** - right panel). To identify tumor-specific TADs among multiple tissues in various groups, we developed a computational algorithm, <u>G</u>roup-, Individual-sample <u>S</u>pecific <u>TA</u>Ds (GISTA), which enables to compare TADs among various groups of tissues as well as at the individual tissue (**METHODS** and **Figure S3B**). At the group-level comparison, GISTA was able to define three categories of TAD changes: inter-tumor conserved TADs (C-TADs), inter-tumor moderately variable TADs (MV-TADs), and inter-tumor significantly variable TADs (SV-TADs) (**METHODS** and **Figure S3D**). At the individual-level comparison, GISTA further subclassified those SV-TADs into two types of individual tumor-specific TADs: highly individual tumor-specific (HS) TADs and lowly individual tumor-specific (LS) TADs. GISTA also identified three special types of individual tumor-specific TAD changes: Neo (N)-Deletion (D), Split (S)-Fuse (F) and Mixed (coexistence of both ND and SF).

**Figure 2.**
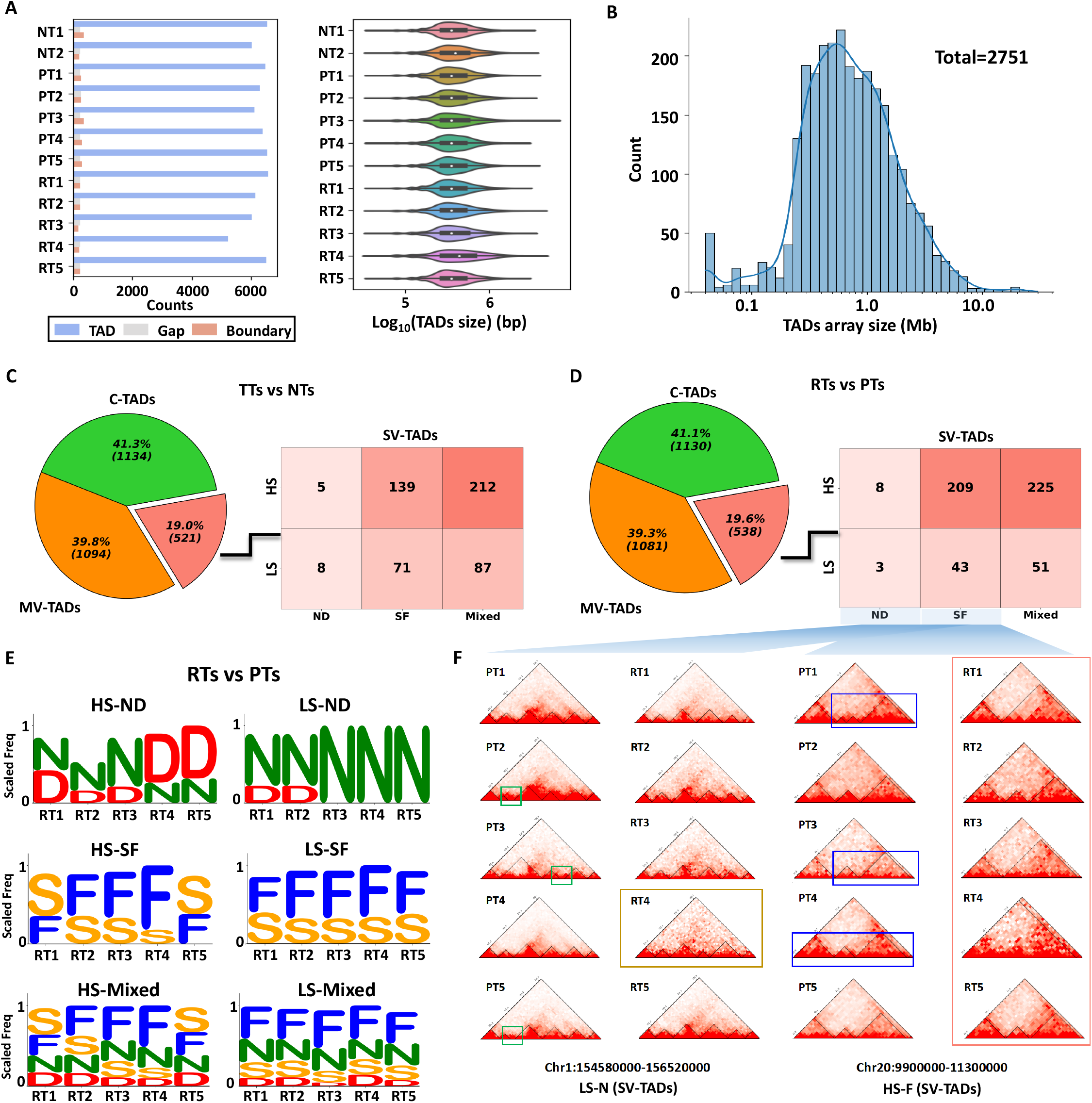
TADs display a highly heterogeneous among breast tumors. **A**. The bar plot (left panel) showing the number of TADs, gaps and boundaries identified in the tissues. The right panel violin plot showing the distribution of TAD’s size. **B**. The histogram displaying the distribution of the size of the TADs array. The total number of TADs array is 2,751. **C**. The pie chart on the left illustrating the proportion of the three comparison types, inter-tumor conserved TADs (C-TADs), inter-tumor moderately variable TADs (MV-TADs), and inter-tumor significantly variable TADs (SV-TADs) between TT *vs* NT. The heatmap on the right demonstrating the number of six sub-categories within SV-TADs, composed of HS-ND, HS-SF, HS-Mixed, LS-ND, LS-SF, and LS-Mixed (**METHODS**). **D**. The pie chart on the left illustrating the proportion of the three types of TAD change between RT vs PT. The heatmap on the right demonstrating the number of six sub-types, HS-ND, HS-SF, HS-Mixed, LS-ND, LS-SF, and LS-Mixed within SV-TADs of RT *vs* PT. **E**. The logo plots showing the scaled frequency of four basic TAD change types, N(eo), D(eletion), S(plit), F(use) for individual RTs by comparing RT *vs* PT in six sub-categories. Top: the frequency of N/D/S/F in HS-ND and LS-ND; middle: the frequency of N/D/S/F in HS-SF and LS-SF; bottom: the frequency of N/D/S/F in HS-Mixed and LS-Mixed. **F**. The visualization of LS-N and HS-F within RT *vs* PT. The green frame indicating ND and SF change, and the pink frame on the left points indicating LS (only RT4 has different TADs array comparing other RTs) and on the right points indicating HS (five RTs have three different TADs array).

After applied GISTA in our pilot, we identified 2,751 TADs arrays with most of the size of arrays ranging from 0.1 to 10 Mb in size (**Figure 2B**). We found that 1,134 (41.3%) of them were C-TADs, 1,094 (39.8%) were MV-TADs, and 521 (19.0%) were SV-TADs between TTs *vs* NTs (**Figure 2C** and **Figure S3E**), whereas 1,130 (41.1%) as C-TADs, 1,081 (39.3%) as MV-TADs, and 538 (19.6%) as SV-TADs between RTs *vs* PTs (**Figure 2D** and **Figure S3E**). Interestingly, we found that the size of the TADs arrays in SV-TADs was larger than those in MV-TADs and C-TADs for both between TTs *vs* NTs and between RTs *vs* PTs (**Figure S4A**). We also observed much more HS type than LS type correlatively with ND, SF and Mixed, with 356 and 166 in TTs *vs* NTs, and 442 and 97 in RTs *vs* PTs, respectively (**Figure 2C, D**). Moreover, Mixed type constituted a majority (>50%) of TAD changes in both TTs *vs* NTs and RTs *vs* PTs comparisons. As expected, we found a more diverse occurrence of N/D/S/F types within HS type for each tissue than within LS type (**Figure 2E** and **Figure S4B, C**). As visualized in **Figure 2F** for two examples, one falling under the LS-N of SV-TADs category on chromosome 1 and the other under the HS-F of SV-TADs category on chromosome 20, we illustrated the TAD changes among the individual tumors. In sum, we identified tumor-specific TAD changes and demonstrated a highly heterogeneous TAD change among breast tumors.

### Chromatin loops show intensively variable among breast tumors

Chromatin looping is a key layer of 3D chromatin architecture that brings distal regulatory elements such as enhancers into proximity with promoters of target genes and has been shown to be linked to tamoxifen resistance in breast cancer cell lines^**13**,**25**^. To identify looping events in tumor tissues and tumor-specific looping genes, we first used FitHiC2^**26**^ to identify a range of ∼100,000-140,000 interacting loci and an average of ∼25,000 promoter (P)-distal (D) loops per tissue, respectively, where an interacting loci was annotated as a P-D loop such that one end of interacting loci is at the promoter region of a UCSC RefSeq gene and the other is within the distal region (200 Kb upstream/downstream to 5’TSS) of the same gene (**Figure 3A** and **Figure S5A**). A gene with at least one P-D loop was further defined as a looping gene. There was an average of ∼12,000 looping genes per tissue (**Figure S5B**). We then calculated a gene-based looping intensity (LI) by combining all P-D loops associated with a particular gene across all 12 tissues (**METHODS** and **Figure S5C**). We then performed a Bi-Gaussian modeling on the Euclidean distance of LI between each of 10 TTs and two NTs (**Figure S5D**) to define TT-specific differential looping genes (DLGs), including Gained (GLGs) or Lost looping genes

**Figure 3.**
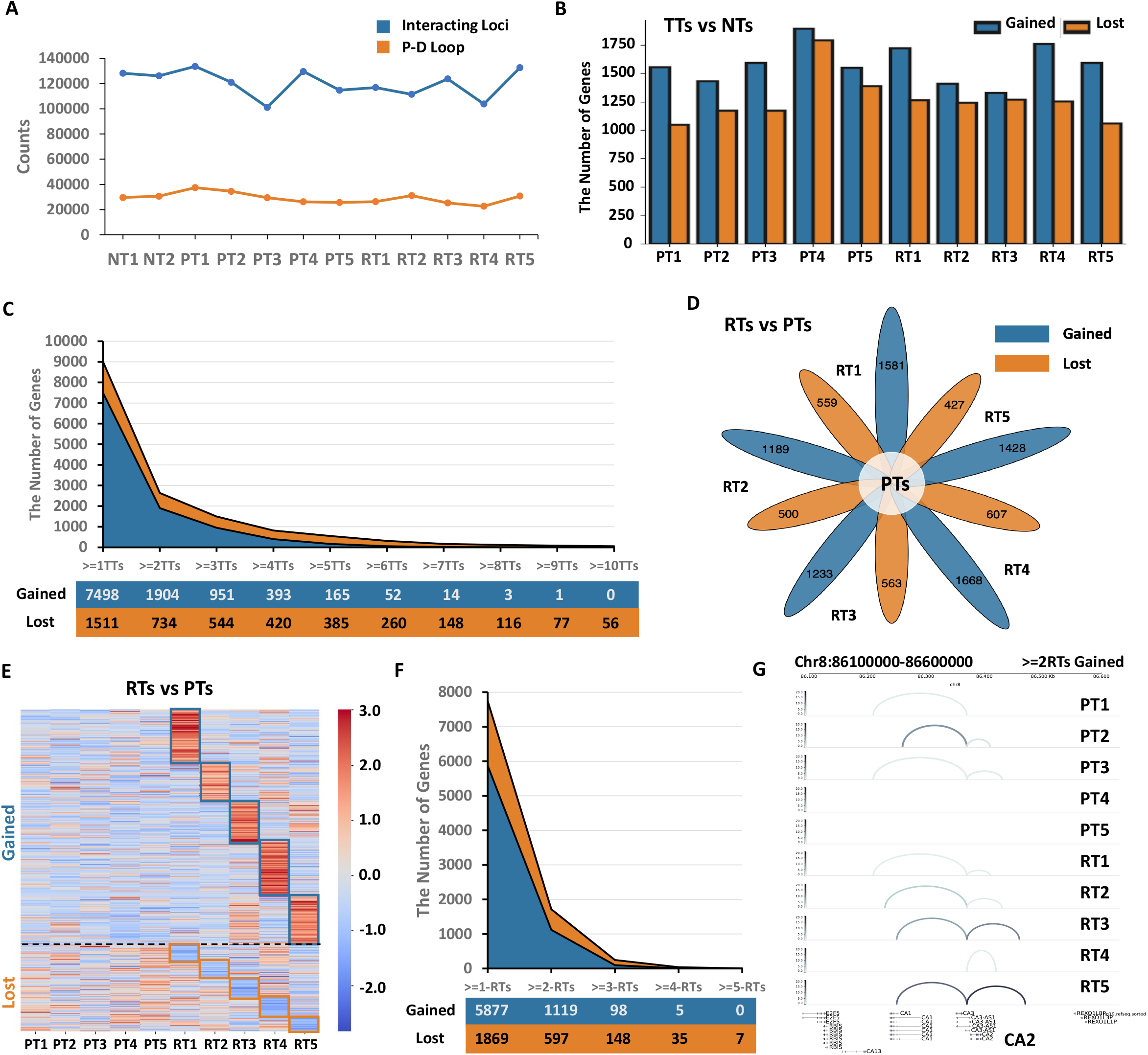
Chromatin loops show intensively variable among breast tumors. **A**. The line plot showing that the number of interacting locus and PD loops (promoter-distal loops for a same gene) of each tissue. **B**. The bar plot illustrating the number of individual tumor-specific looping genes. **C**. The stacked line plot showing the number of common DLGs among the 10 TTs. **D**. The flower plot showing the number of individual RT-specific looping genes compared to PTs. **E**. The heatmap illustrating the loop intensity of individual RT-specific DLGs. **F**. The stacked line plot showing the number of common DLGs among the five RTs. **G**. The genome tracks displaying the CA2 DL loops, belonging to >=2RTs GLGs. The color bar representing the intensity of loops ranging from 0 to 20. The darker the color, the higher the intensity of loops.

(LGLs) based on the sign of the distance. We identified a range of 1,326-1,895 Gained and a range of 1,049-1,792 Lost looping genes for each of 10 TTs, respectively (**Figure 3B** and **Figure S5E**). Remarkably, we observed a negative correlative trend that the number of shared Gained or Lost looping genes was decreasing along with the increase of the number of shared tumors (**Figure 3C**), suggesting looping genes are quite a diverse among individual tumors. In parallel, we identified a range of 1,189-1,668 Gained and a range of 427-607 Lost (**Figure 3D**) looping genes (**Figure 3E**) for each of five RTs, respectively. We also observed a similar trend that the number of shared Gained or Lost looping genes was decreasing along with the increase of the number of shared RTs (**Figure 3F** and **Table S3**). Bird views of a few Gained looping genes, CA2, NUBPL, and ATP2B2 were showed in **Figure 3G** and **Figure S5F**, respectively. Collectively, our results revealed intensively variable chromatin looping events among individual breast tumors, suggesting a looping-mediated tumor heterogeneity during the progression to endocrine resistance.

### Cross-examination identifies RT-specific looping-mediated biological pathways

We further conducted a qualitative comparison among the set of RT-specific differential looping genes (5,877 gained and 1,869 lost), four differential compartments (A-A, A-B, B-A, and B-B) and six types of SV-TADs (HS-ND, HS-SF, HS-Mixed, LS-ND, LS-SF, and LS-Mixed) (**Figure 4A**). We observed that the number of looping genes associated with differential compartments was substantially lower than those associated with TAD changes. This might primarily be due to the fact that compartment segments at a 100kb bin-size resolution instead of the whole compartment were considered as differential by dcHiC. Interestingly, a positive correlation was identified between individual HS TADs and differential looping genes (**Figure 4B**). To explore a correlation of the three layers of chromatin architecture with copy number variations (CNVs), we further performed an analysis by applying HiNT^**27**^ on tissue Hi-C data and identified numerous CNVs, including copy number gain, loss, and neutral variants (**Figure S6A-C**). We discovered a higher percentage of overlapping between compartments and copy number gain/loss in tumor tissues (**Figure S6D**). Additionally, we also observed that the contact frequency within TAD boundaries around the CNV breakpoints was notably low^**16**^ (**Figure S6E**), and the total number of loops located in copy number gain/loss regions was greater in RTs than in PTs (**Figure S6F**).

**Figure 4.**
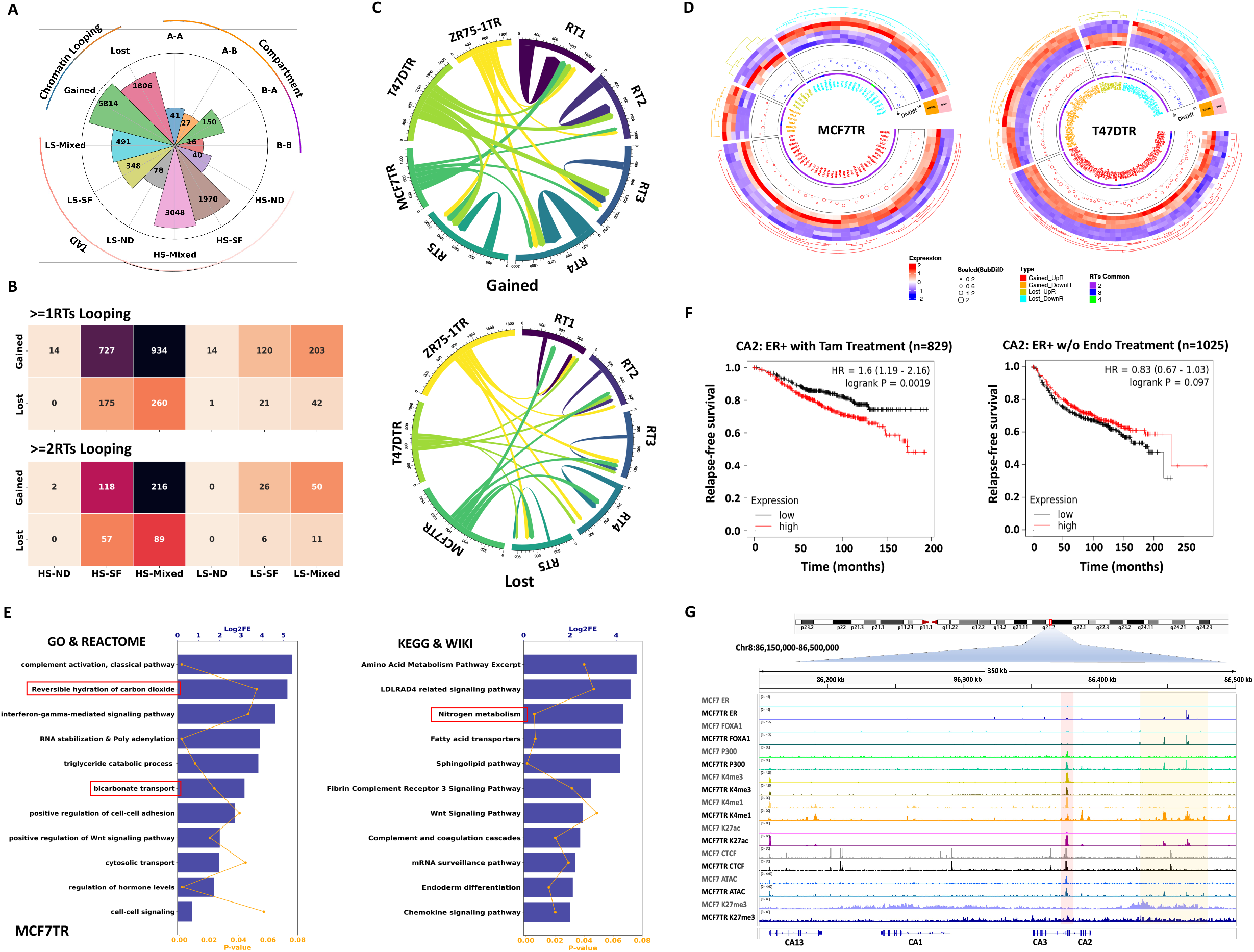
Cross-examination identifies RT-specific looping-mediated biological pathways. **A**. The circle bar plot showing the number of common genes among the changes of three layers of chromatin architecture. **B**. The heatmaps showing the number of common genes between six types of TAD change and two types of looping change. The top panel was for at least one IDLG while the bottom panel was for at least two IDLGs. **C**. The top chord diagram showing the overlapping relationship of GLGs between five RTs and three TR cell lines (MCF7TR, T47DTR and ZR75-1TR). Among three TR cell lines, MCF7TR has the highest correlation (216 GLGs) with RT1, T47DTR has the most overlapping with RT2 to RT5 (204, 225, 228 and 235 GLGs, respectively), and ZR75-1TR has the relatively mild correlations with all RTs compared to MCF7TR and T47DTR. The bottom chord diagram showing the overlapping relationship of LLGs between five RTs and three TR cell lines. Among three TR cell lines, MCF7TR has the highest correlation with most of the RTs: RT1, RT2, RT3 and RT5 (125, 85, 122 and 99 LLGs, respectively), T47DTR has the relatively mild correlations with all RTs compared to MCF7TR and ZR75-1TR, and ZR75-1TR has the most overlapping with RT3. **D**. Circular heatmap-bubble plot showing all >=2RTs_MCF7TR DELGs (top) and >=2RTs_T47DTR DELGs (bottom) with four properties: mRNA expression, loop intensity difference (DivDiff and SubDiff), the number of common RTs. The outermost heatmap showing the scaled gene expression of four types of DELGs with four dendrogram colors: red, orange, yellow and cyan for Gained/UpR, Gained/DownR, Lost/UpR, Lost/DownR, respectively; The middle lay bubble plot showing the scaled DivDiff: blue for the gene has scaled DivDiff value< 0 and red for the gene has scaled DivDiff value> 0 as well as scaled SubDiff: the larger of the circle size, the larger the scaled SubDiff value; and the innermost layer indicates the number of common RTs of a certain DLG. **E**. Bar plots showing the top enriched terms of GO, REACTOME, KEGG and WIKI for >=2RTs_MCF7TR DELGs. The pathways within the red frames is composed of CA1 and CA2. **F**. Kaplan-Meier plots of CA2 showing the probability of relapse-free survival in ER+ breast cancer patients with tamoxifen treatment (n = 829) and without endocrine treatment (n = 1025). CA2 gene expression was classified as low or high (black or red lines, respectively) based on the comparison of its median cut-off value. p value was determined by the log-rank test. **G**. The IGV visualization of the enrichment of ER, FOXA1, P300, CTCF, H3K4me3, H3K4me1, H3K27ac, H3K27me3, and ATAC on the promoter and enhancer regions of CA2.

We next cross-examined the tumor-specific DLGs with the cell-based DLGs. We first used the same method of analyzing tumor-based Hi-C data to re-analyze the Hi-C data on the three breast cancer cell systems, MCF7 *vs* MCF7TR, T47D *vs* T47DTR and ZR75-1 *vs* ZR75-1TR from our previous study^**28**^. We thus obtained TR-specific DLGs for each of the three cell systems. We then overlapped each of the three TR-specific DLGs with the set of >=1RTs RT-specific DLGs and found the common looping genes among different sets were quite diverse (**Figure 4C**). We further derived the differentially expressed looping genes (DELGs) by correlating common DLGs with differential expressed genes (DEGs) for MCF7/MCF7TR and T47D/T47DTR cells respectively. We identified a total of 68 DELGs, including 35 Gained/UpR, 21 Lost/DownR, 6 Gained/DownR and 6 Lost/UpR for MCF7/MCF7TR as well as a total of 115 DELGs, including 54 Gained/UpR, 21 Lost/DownR, 31 Gained/DownR and 9 Lost/UpR (**Figure 4D**). Furthermore, we performed the gene ontology (GO) and biological pathway analyses by using DAVID^**29**^ and Enrichr^**30**^. For MCF7TR-specific DELGs, we found reversible hydration of carbon dioxide, interferon gamma mediated signaling pathway, and bicarbonate transport were among the top enriched pathways under GO and REACTOME analyses whereas amino acid metabolism, nitrogen metabolism and Wnt signaling pathway under KEGG and WIKI analyses **(Figure 4E**). The results of GO, REACTOME, KEGG and WIKI analyses on T47DTR-specific DLGs were showed in **Figure S7A**. Surprisingly, there were very few common pathways between two cell systems, including RNA stabilization, bicarbonate transport and nitrogen metabolism, in which these pathways encompass nine MCF7TR-specific DELGs, AXIN2, CA1, CA2, C4BPA, C4BPB, CBLB, DDO, DTD2 and IL7R, and seven T47DTR-specific DELGs, AQP1, CA1, CA2, COX6C, HMGCS1, TP63, and UQCR10, respectively. Since CA1 and CA2 are common between MCF7TR and T47DTR, we subsequently performed survival analyses^**31**^ on these two genes and found that high expression levels of both genes were associated with a worse relapse-free survival in breast cancer patients treated with tamoxifen (**Figure 4F** and **Figure S7B**). We examined the CA2 distal loop with the enrichment of various transcription factors (TFs) and histone marks, and found a higher enrichment of ER, FOXA1, P300, H3K4me3, H3K4me1, H3K27ac, CTCF and ATAC-seq signals but lower enrichment of H3K27me3 in MCF7TR *vs* MCF7 cells (**Figure 4G**), indicating that an enhancer-promoter looping (EPL)-mediated CA2 might contribute to the tamoxifen resistance. Together, our analysis identified RT-specific looping-mediated biological pathways by cross-examining tumor-based and cell-based differential looping genes.

### The inhibition of CA2 impedes tumor growth both *in vitro* and *in vivo*

Numerous previous studies have demonstrated that CA2 played an important role in tumor progression, metastasis and treatment resistance^**32-36**^. Our comprehensive analysis not only suggested a role of EPL-mediated CA2 in contributing to the tamoxifen resistance, but also pinpointed it as a promising therapeutic target gene. Brinzolamide, a commercially available inhibitor targeting CA2, has been tested in its antitumor activities in Glioblastoma and blood cancer^**37**,**38**^. We thus attempted to test its inhibition efficacy in tamoxifen resistant breast cancer cells by using both *in vitro* phenotypical assays and *in vivo* cell-derived xenograft (CDX) mice model.

We first applied the CCK-8 assay to evaluate the effect of Brinzolamide on MCF7/MCF7TR and T47D/T47DTR cell growth and to determine their respective optimal inhibitor concentrations over a 0 to 6-day period (**Figure S8A, B**). We observed a stronger inhibition of cell growth or impeding of cell proliferation in both TR cells than in two Tam-sensitive (TS) cells, MCF7 and T47D, respectively (**Figure 5A, B**). We then tested Brinzolamide’s therapeutic efficacy in CDXs by utilizing MCF7 and MCF7TR cells in estrogen disk-implanted female nu/nu mice. Following the development of palpable tumors, the mice were administered either Brinzolamide or a control solution for a five-week period. Notably, the MCF7TR CDXs in the Brinzolamide-treated group exhibited a significantly slower tumor growth rate compared to those in the control group (**Figure 5C**) whereas MCF7 CDXs did not show any significantly decrease in tumor growth rate (**Figure S8C**). Crucially, throughout the treatment and post-treatment periods, the mice showed no signs of drug-induced toxicity, and there were no substantial weight differences between the treated and control groups for both CDX models (**Figure 5D** and **Figure S8D**). After the mice reached the predetermined tumor burden as per the protocol, all mice were euthanized, and the mammary tumors were excised. Tumors from Brinzolamide-treated MCF7TR CDXs were significantly smaller than those from the control group, a pattern not seen in MCF7 CDXs (**Figure 5E** and **Figure S8E**). Our findings from both *in vitro* and *in vivo* functional characterization underscore the therapeutic potential of targeting CA2 in treating tamoxifen-resistant breast cancer.

**Figure 5.**
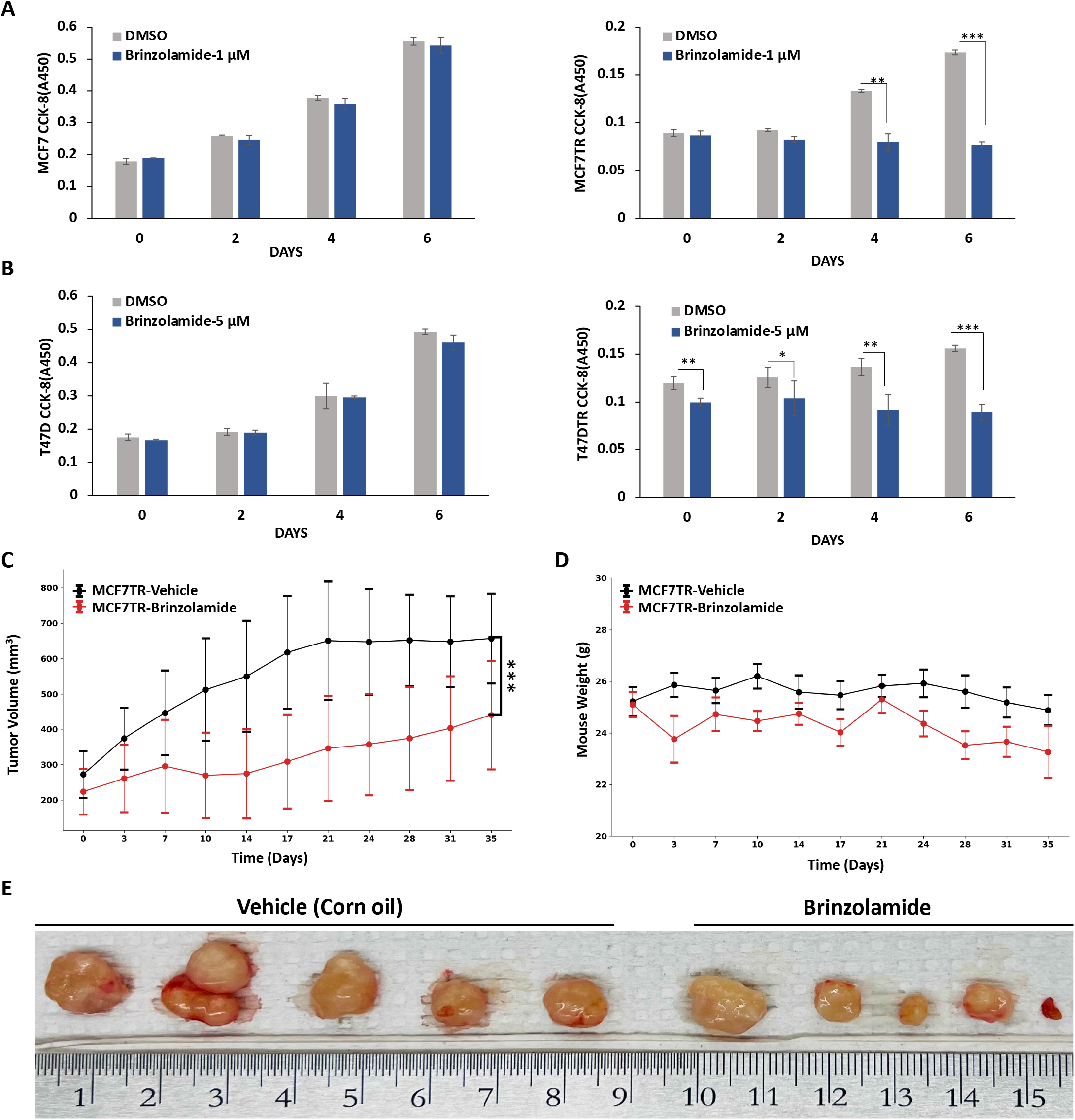
The inhibition of CA2 impedes tumor growth both *in vitro* and *in vivo*. **A**. CCK-8 assay was used to detect the cell proliferation in MCF7 and MCF7TR cells seeded at 1000 cells per well in a 96-well plate. MCF7 and MCF7TR cells were treated with Brinzolamide (1μM). Values were expressed as the mean□±□Standard deviation of three independent experiments. Samples t-test **p < 0.005, ***p < 0.001 was considered statistically significant. **B**. CCK-8 assay was used to detect the cell proliferation in T47D and T47DTR cells seeded at 1000 cells per well in a 96-well plate. T47D and T47DTR cells were treated with Brinzolamide (5μM). Values were expressed as the mean□±□Standard deviation of three independent experiments. Samples t-test *p < 0.05, ** p < 0.005, ***p < 0.001 was considered statistically significant. **C**. Brinzolamide inhibited *in vivo* tumor growth of MCF7TR xenograft mice. MCF7TR cells were inoculated into the mammary fat pads of nude mice. When tumors became palpable, tumors were treated with Vehicle (Corn oil) and Brinzolamide, respectively. Tumor growth was analyzed by measuring the tumor volume. Tumor volume (length x width) over time for each treatment group, monitored via caliper measurements. Error bars represent standard error mean. Significance is shown only for endpoint measurements. Samples t-test ***p < 0.001 was considered statistically significant. **D**. Mice weights over time for Brinzolamide-treated vs Vehicle group. Error bars represent the standard error mean. **E**. Photographs of mammary tumors for Brinzolamide-treated *vs* Vehicle groups at the study endpoint. The ruler scale is mm.

### The inhibition of CA2 reverses chromatin looping

To assess whether the inhibition of CA2 might alter the chromatin looping and gene expression of CA2 itself as well as other looping-mediated TR-specific genes within the transport and metabolism pathways, we conducted 3C-qPCR and RT-qPCR analyses on the selected genes in MCF7/MCF7TR and T47D/T47DTR cells. Upon the treatment of Brinzolamide, we observed a significantly decreased P-D looping interaction of CA2 in both MCF7TR and T47DTR cells, in contrast to their respective TS cells (**Figure 6A, B**). Additionally, Brinzolamide led to diminished P-D looping interactions for other looping genes including CA1, AXIN2, C4BPA, CBLB, IL7R, DDO, and DTD2 in MCF7TR cells, and CA1, COX6C, TP63, and HMGCS1 in T47DTR cells (**Figure 6A, B**). These results indicated that Brinzolamide was able to reverse chromatin looping activities. The gene expression level showed a reversed expression of CA2 between MCF7 and MCF7TR, as well as between T47D and T47DTR cells, respectively (**Figure 6C, D**). However, this reversal expression was not so evident for some other genes in the TR cells. Nevertheless, our data revealed a mechanistic link between looping-mediated CA2 and the transport and metabolism pathways in promoting breast cancer cell tamoxifen resistance. In summary, the silence of CA2 through a pharmacological drug inhibition is able to reverse the chromatin looping of CA2 and some other RT-specific looping genes, suggesting CA2 might play an oncogenic role in driving the progression of breast cancer endocrine resistance through a chromatin looping mechanism.

**Figure 6.**
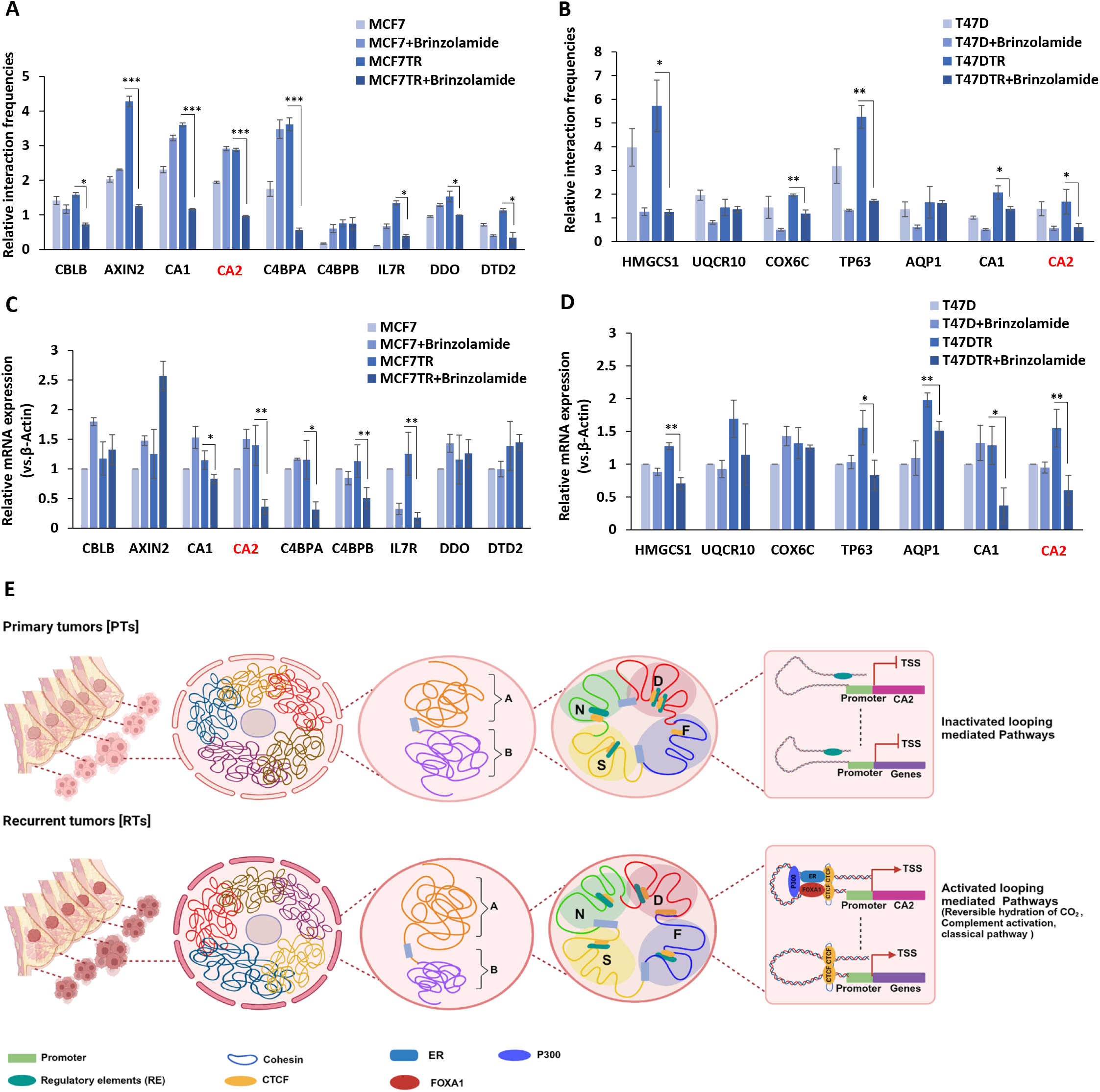
The inhibition of CA2 reverses chromatin looping. **A**. The relative interaction frequencies of the selected genes in MCF7 and MCF7TR cells with and without Brinzolamide identified by 3C-qPCR. Values were expressed as the mean□±□Standard deviation of three independent biological replicates *p < 0.05, ** p < 0.005, ***p < 0.001. **B**. The relative interaction frequencies of the selected genes in T47D and T47DTR cells with and without Brinzolamide identified by 3C-qPCR experiments. Values were expressed as the mean□±□Standard deviation of three independent biological replicates *p < 0.05, ** p < 0.005. **C**. Quantitative Real-time PCR was performed to detect the mRNA expression level of the selected genes in MCF7 and MCF7TR cells with and without Brinzolamide. Values were expressed as the mean□±□Standard deviation of three independent biological replicates *p < 0.05, ** p < 0.005. **D**. Quantitative Real-time PCR was performed to detect the mRNA expression level of the selected genes in T47D and T47DTR cells with and without Brinzolamide. Values were expressed as the mean□±□Standard deviation of three independent biological replicates *p < 0.05, ** p < 0.005. **E**. A possible looping-mediated mechanistical model illustrating that inactivated biological pathways due to the weaken E-P looping in PTs became activated by enhanced looping activities in RTs, underscoring a role of chromatin looping in regulating downstream biological pathways and further driving the breast tumor heterogeneity of endocrine resistance.

## DISCUSSION

Despite many studies have linked the alterations of 3D chromatin organization to the cancer development and treatment resistance^**15**,**28**,**39-43**^, including breast cancer endocrine resistance^**15**,**41**,**42**^, most of these studies were conducted in cancer cell culture models except a few very recent studies conducted in human tumor samples^**17-20**^. Those Hi-C studies in tumor samples clearly revealed much sophisticated 3D genome regulatory pattern including inter- and intra-tumor heterogeneity. To this end, we designed this study to exploit the 3D chromatin variation in a pilot of 12 breast normal and tumor tissues. Our comprehensive analysis of the three layers of chromatin architecture uncovered inter-tumor heterogeneities among breast tumors in all of three layers, compartment, TAD and chromatin looping. Furthermore, our cross-examination of tumor-based looping genes and cell-based looping genes identified the common looping genes enriched with the transport and nitrogen metabolism pathway. Of the gained looping and upregulated genes within this metabolism pathway, CA2 showed a worse relapse-free survival in breast cancer patients treated with tamoxifen (**Figure 4F**), indicating its potential oncogenic role in driving the progression of breast cancer endocrine resistance. Indeed, a pharmacological drug inhibition to CA2 impeded the tumor growth both *in vitro* and *in vivo* (**Figure 5**). Our study thus advances a novel concept that endocrine-treatment responses could be governed by individual tumor-specific 3D chromatin domains which clearly departs from the prevailing paradigm that endocrine-treatment responses are driven by the genetic variability between individual patients. Therefore, this work will yield significant mechanistic insights into the role and clinical relevance of 3D chromatin architecture in breast cancer endocrine resistance.

Our data demonstrated a conserved compartmentalization in terms of the size and the number of compartments A and B across breast tumors, a finding consistent with other studies in leukemia and colon cancer^**17**,**18**^. Our data also suggested that chromatin architecture is largely preserved at the compartment layer. However, TADs, another layer of chromatin architecture, display a highly heterogeneous among breast tumors. Since no computational tools are currently available to identify TADs among the group of tumor samples and for the individual tumor sample, we developed GISTA, a novel algorithm specifically designed to identify significantly variable TADs among different groups of multiple samples. The novelty of GISTA includes 1) comparing the difference between two groups based on a series of TADs arrays instead of individual TAD; 2) applying a dynamic programming to capture the boundary of TADs array with a user-defined boundary-shift parameter; and 3) utilizing a decision tree to annotate different types of TAD changes. Indeed, our tool was able to identify many different types of TAD changes, including conserved and individual PT- or RT-specific TADs, demonstrating its high applicability in two or three group of tumors.

While it has been documented that breast cancer cell-specific chromatin looping play a crucial role in regulating the expression of oncogenes^**44**,**45**^, a comprehensive understanding of the chromatin looping variability in tamoxifen resistance is still lacking. Remarkably, we observed the common differential looping genes (both gained and lost) among tumors decrease with the increase of the number of shared tumors (**Figure 3C**). A similar trend was also observed in recurrent tumors (**Figure 3F**). The extensive variabilities in chromatin looping across tumors, especially in recurrent tumors, highlight an important role of chromatin looping in regulating breast tumor heterogeneity. We also found a positive correlation of differential looping genes with highly individual tumor-specific (HS) TADs but not with differential compartments, in which it could be possibly explained by the model of TAD formed through a loop extrusion^**46-48**^, whereas compartments are formed by a distinct mechanism^**49**^. Despite other studies suggested the variations in the intrinsic chromatin interaction landscape were associated with differential gene expression^**50**^, our data demonstrated a low correlation between differential chromatin looping genes with differential gene expression. Nevertheless, from the set of RT-specific differentially expressed looping genes, we were able to identify several interesting signaling pathways including RNA stabilization, bicarbonate transport and nitrogen metabolism which might be involved in regulating tamoxifen resistance.

Notably, we identified CA2, an enzyme of the family of carbonic anhydrases, as a potential driver in the progression of tamoxifen resistance. Carbonic anhydrases (CAs) facilitate the conversion of CO2 and H2O into bicarbonate, playing vital roles in diverse cellular processes^**37**,**51**^. Although previous work has linked the expression of CA2 with the patient survival rates in Luminal B breast cancer^**52**^, its role in tamoxifen resistant breast cancer remains unexplored. Our functional examination demonstrated that CA2 inhibition impeded cell proliferation in MCF7TR and T47DTR cells as well as slowed tumor growth in xenograft mouse models upon the treatment of a CA2-specific inhibitor, Brinzolamide^**53-55**^. Furthermore, we showed a reversable chromatin looping and gene expression of CA2 and other genes in TR cells after CA2 inhibition, suggesting CA2 might play an oncogenic role in driving the tamoxifen resistance through a chromatin looping mechanism. Nevertheless, additional experiments are needed to elucidate the underlying mechanism of how looping-mediated CA2 in regulating the downstream biological pathways and further driving the tamoxifen resistance.

Collectively, we propose a possible looping-mediated mechanistic model for driving the breast cancer tamoxifen resistance: in primary tumors or tamoxifen-sensitive cells, the looping-mediated pathways and their component genes are inactive due to their weakened enhancer-promoter (E-P) interactions, while in tamoxifen-treated recurrent tumors or tamoxifen-resistant cells, these E-P interactions are strengthened, leading to activation of genes and pathways (**Figure 6E**). Our model is further substantiated by the following observation: increased chromatin accessibility and active marks, H3K4me1 and H3K27ac, strengthened ER, FOXA1, P300 and CTCF bindings whereas a decreased repressive mark, H3K27me3 in the CA2 enhancer region in MCF7TR *vs* MCF7 cells (**Figure 4G** and **Figure 6E**). We thus hypothesize that CA2 is silenced by loss of ER signaling^**56**^ in Tam-sensitive breast cancer cells. A prolonged Tam-treatment mediates a non-genomic calcium signaling resulting in activation of kinase pathways such as MAPK/ERK which can modulate ER membrane translocate to the nucleus and promote CA2 activation through a looping regulation. This non-genomic and genomic crosstalk can in turn enhance cell migration and proliferation^**57**^ resulting in Tam-resistance. Future experiments will test this hypothesis with various functional and molecular assays, such as CRISPR/Cas9, 3D-FISH, 3C/ChIP/RT-qPCR.

In summary, our study has produced a rich resource of high-quality 3D chromatin data in a pilot cohort of tumor tissues, in which they can further be used for identifying 3D chromatin guided prognostic biomarkers to stratify breast cancer that may acquire resistance to endocrine therapy, and subsequently designing more effective therapeutic strategies to overcome tamoxifen resistance. To the best of our knowledge, this is the first investigation of 3D chromatin variabilities in breast tumor tissues. The tumor-specific 3D chromatin alterations provide a significant mechanistic insight into the role of 3D chromatin-regulated tumor heterogeneity in breast cancer endocrine resistance.

## STAR⍰METHODS

Detailed methods are provided in the online version of this paper and include the following:

- **KEY RESOURCES TABLE**
- **RESOURCE AVAILABILITY**
  ∘ Lead contact
  ∘ Materials availability
  ∘ Data and code availability
- **EXPERIMENTAL MODEL AND STUDY PARTICIPANT DETAILS**
  ∘ **Animal care**
- **METHOD DETAILS**
- **QUANTIFICATION AND STATISTICAL ANALYSIS**

## SUPPLEMENTAL INFORMATION

Supplemental information can be found online at https://doi.org/.

## ACKNOWLEDGMENTS

We thank the UTHSA Next Generation Sequencing Facilities, Dr. Zhao Lai for sequencing Hi-C data. This project was partially supported by grants from NIH R01GM114142 and Advancing A Healthier Wisconsin (AHW) Seed Grant.

## AUTHOR CONTRIBUTIONS

VXJ conceived the project. LC conducted the experiments. KF performed the data analyses. TL assisted the data analysis. VXJ, LC, KF, and TL wrote the manuscript.

## STAR⍰METHODS

### KEY RESOURCES TABLE

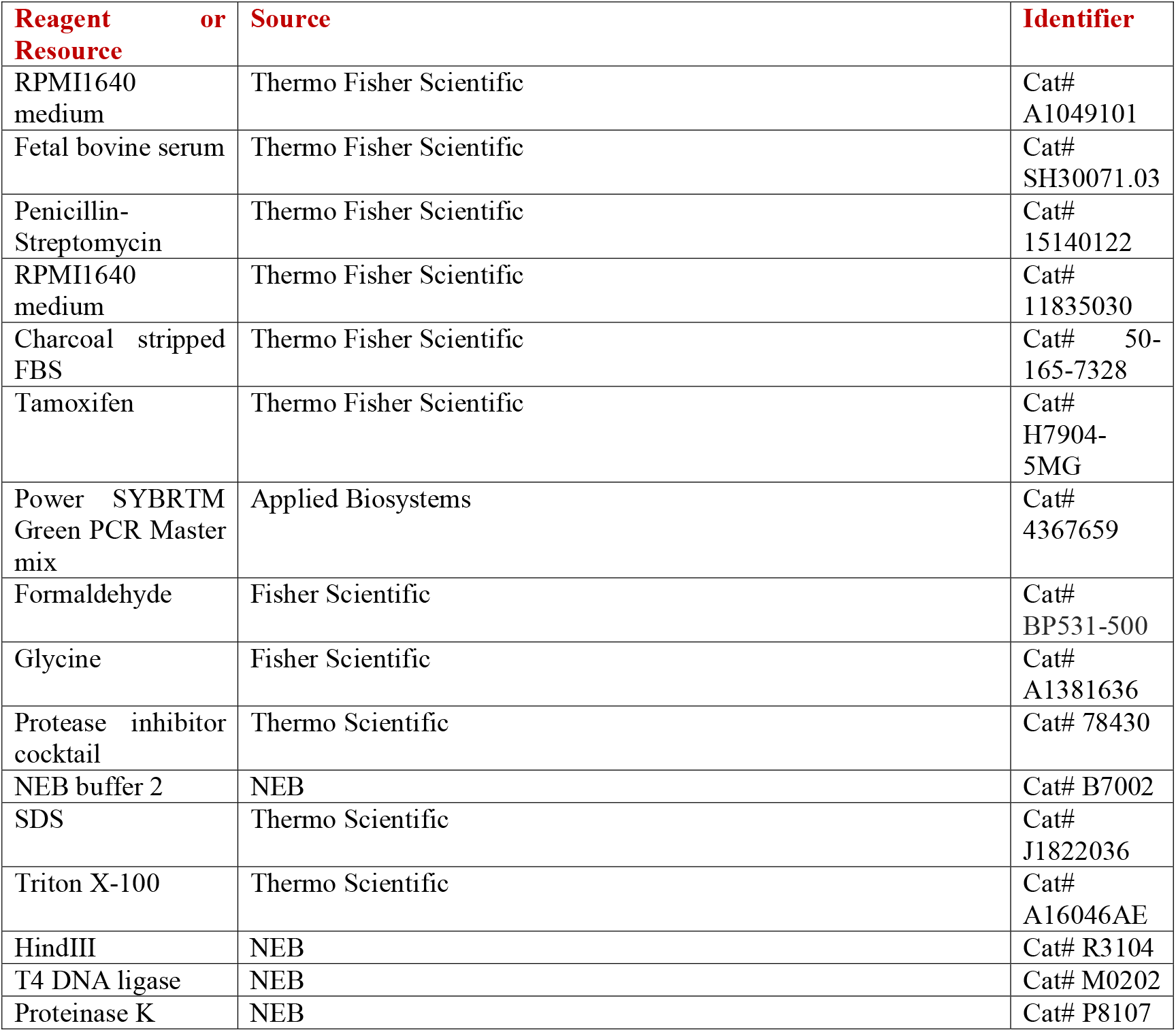
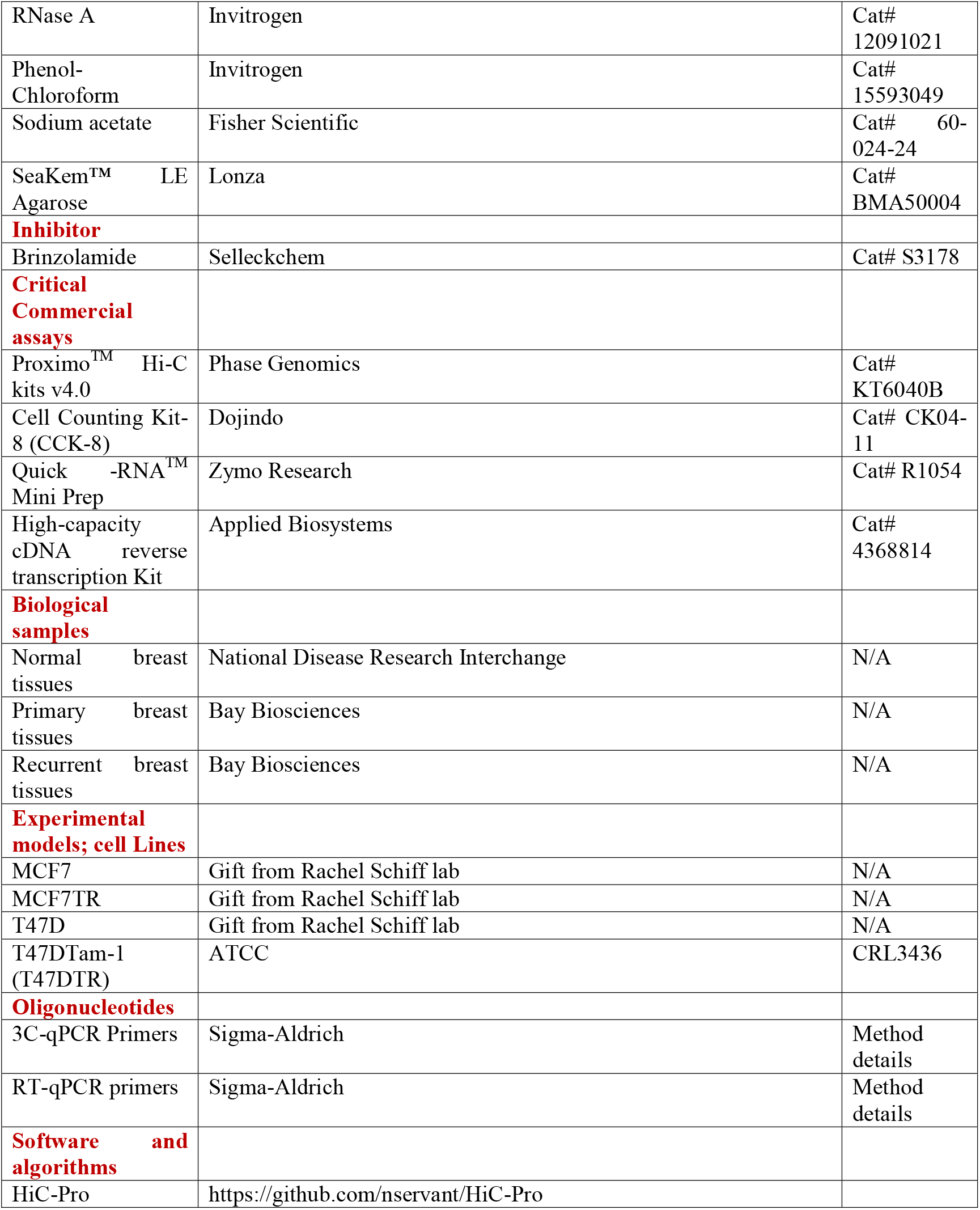
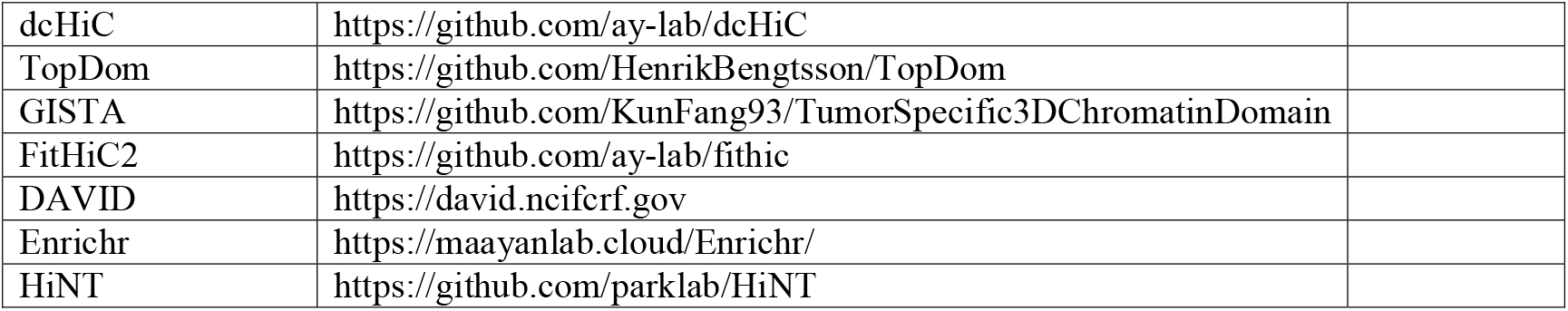

## RESOURCE AVAILABILITY

### Lead contact

Further information and requests for resources should be directed to and will be fulfilled by the lead contact, Victor Jin.

### Materials availability

This study did not generate new unique reagents.

### Data and code availability

Raw and processed Hi-C data for tissue samples were deposited in GEO under accession number GSE261230. Other public datasets analyzed during the current study are available in the GEO repository: GSE144380^**28**^, GSE108787^**15**^, GSE128676^**28**^, and GSE128460^**58**^.

The source codes of this study are freely available at Under GNU General Public License (GPL-v3.0) (https://github.com/KunFang93/TumorSpecific3DChromatinDomain).

## EXPERIMENTAL MODEL AND STUDY PARTICIPANT DETAILS

Five fresh primary and five fresh tamoxifen-treated recurrent breast tumor samples were purchased from Bay Biosciences LLC (Brookline, MA). Two normal breast tissues were procured from National Disease Research Interchange (NDRI, Philadelphia) as fresh frozen pieces (**Table S1**). All the tissues were stored at -80°C until they were used for experiments. The use of breast normal and tumor tissues for this study was conducted under MCW approved IRB approval.

## METHOD DETAILS

### Hi-C profiling on breast tissues

Approximately 100-200 mg of each tissue was chopped into fine pieces before being fixed. Hi-C was performed using Phase Genomics Proximo Hi-C kits v4.0 (Phase genomics, Seattle, WA) according to the manufacturer’s instructions^**21**^. Briefly, the finely chopped tissues were incubated with crosslinking solution for 15 minutes with gentle rotation. Tissue lysis was performed with motor and pestle with liquid nitrogen and grinded to a fine powder, and resuspended in 700 μl of Lysis buffer 1. The chromatin containing supernatant was pelleted and washed with 500 μl of 1x CRB, resuspended in 100 μl of Lysis Buffer 2. 100 μl of SPRI beads were added, incubated and washed with 200 μl of 1x CRB. SPRI beads containing chromatin were resuspended in 150 μl of fragmentation buffer, 5 μl of restriction enzyme (Sau3AI) and incubated at 37 °C for 1 hour. 2.5 μl of finishing enzyme is added and mixed by vortexing. Ligation was performed by adding 95 μl of Ligation buffer and 5 μl ligation enzyme to the SPRI beads containing chromatin. To revert the crosslinking, 5 μl of RX enzyme was added to the reaction and then incubated overnight at 65 °C with shaking (900 rpm). The pellet was washed, air dried, resuspended in 100 μl of elution buffer and then incubated for 5 minutes at room temperature. The beads were washed and resuspended in 100 μl of bead binding buffer. After the final wash the beads were resuspended in 200 μl of deionized water and measured the concentration of DNA using Qubit dsDNA HS assay kit. About 500 ng of streptavidin beads containing DNA were separated, resuspended in 20 μl of deionized water, and a total of 25 μl of library reagent and an approximate amount of library reagent 1 is added. The reaction was mixed gently and incubated at 55 °C for 10 min. PCR was performed by adding 26 μl of HSR mix (PCR Hotstart Ready mix) and 5 μl of PCR index primer to each reaction. After the amplification, the sample was size selected by adding 25 μl of SPRI beads to the library. The beads were separated and washed twice with 200 μl of 80% ethanol for at least 30 sec each and the pellet was air dried for no longer than 5 min at room temperature. The beads were resuspended and separated and the supernatant containing the final Hi-C library which was quantified using Qubit dsDNA HS assay kit on a Qubit. The library was sequenced on Illumina HiSeq3000.

### Cell lines and reagents

Human breast cancer cell lines MCF7, T47D and their tamoxifen resistant (TR) cells were derived from previous studies^**59-62**^. Both MCF7 and T47D cells were cultured in RPMI1640 medium (Thermo Fisher Scientific, Catalog #A1049101) supplemented with 10% fetal bovine serum (FBS) (Thermo Fisher Scientific, Catalog # SH30071.03) and 1% Penicillin-Streptomycin (Thermo Fisher Scientific, Catalog # 15140122). Tamoxifen resistant cells were cultured in phenol-red free RPMI1640 medium (Thermo Fisher Scientific, Catalog # 11835030) with 10% charcoal-stripped FBS (Thermo Fisher Scientific, Catalog # 50-165-7328) and 1% Penicillin-Streptomycin and 100nM tamoxifen (Sigma-Aldrich, Catalog #H7904-5MG). Tamoxifen was replaced every 48h. All the cells were grown at 37°C and 5% CO2 until they reach 90% confluence. The cells were treated with Brinzolamide (Selleckchem, Catalog # S3178). The gradient concentrations of Brinzolamide are 1 μM for MCF7 cells and 5 μM for T47D cells respectively. Cell absorbance was recorded at different time points.

### CCK-8 cell viability assay

Cell viability was measured by CCK-8 (CCK-8, Dojindo, USA) assay following the manufacturer’s instruction. In brief, MCF7, T47D and their tamoxifen resistant (TR) cells were harvested and plated at a density of 1 x 10^3^ cells per well in 96-well plates (Corning Inc) and cultured in an incubator5% CO2 incubator at 37 °C. After 24 h the culture media was replaced, and the cells are incubated with various concentrations of Brinzolamide. At the end of each time point 10 μL of CCK-8 solution was added to each 96-well plates and the mixture was incubated for 1 hour in the incubator at 37 °C. The optical density at 450 nm was measured at different time points using BioTek ELx800 Absorbance Microplate Reader^**28**^. The experiments were repeated and analyzed three times separately.

### RNA isolation and RT-qPCR

Total RNA was isolated using Quick -RNATM Mini Prep (Zymo Research, USA) according to the manufacturer’s instructions. Five million cells from MCF7 and MCF7TR were lysed in RNA lysis buffer followed by eliminating the majority of gDNA with Spin-Away Filter. Then the mixture of RNA and ethanol was loaded onto Zymo-Spin IIICG Column followed by DNase I treatment to remove the traces of DNA. The samples are washed twice with RNA wash buffer and the total RNA was eluted in 50μL DNase/RNase-Free Water. Two microgram of RNA was used to convert to cDNA using a high-capacity cDNA reverse transcription Kit (Applied Biosystem). qPCR was conducted with a Power SYBRTM Green PCR Master mix (Applied Biosystem). Real-time qPCR was performed on QuantStudio 3 Real-Time PCR system (Applied Biosystem, USA) according to the manufacturer’s instructions. The relative expression of RNAs was determined by the ΔΔCT method using ACTB as an internal control for quantification analyses of gene targets^**28**,**58**^. Primers used are listed in (**Table S4**). Each PCR reaction was performed in triplicate, and the data presented were the average of three independent experiment results for all PCR reactions.

### 3C-qPCR

3C-qPCR experiment was performed as previously described^**63**^. Approximately five to ten million cells of MCF7, T47D and their tamoxifen resistant (TR) cells were cross-linked with 1% formaldehyde for 10 min at room temperature. The reaction was quenched by 1 M glycine for 5 min at room temperature. Cells were lysed with 500μl of cold lysis buffer (10 MmTris–HCl Ph 8.0, 10 Mm NaCl, 0.2% Igepal CA630) with protease inhibitors for 1 h on ice. After lysis, the cell nuclei were pelleted, and the chromatin was digested using 200 units of HindIII (NEB) at 37 °C overnight and then the digestion was stopped with 1.6% SDS at 65 °C for 20 min. Digested DNA fragments were ligated using T4 DNA ligase (NEB) for 4 h at 16 °C. Samples were reverse cross-linked with Proteinase K at 65 °C overnight. 3C samples were then purified using phenol– chloroform extraction. The 3C template was dissolved in 10 mM Tris-HCl and DNA concentrations were measured using Nanodrop. For3C several primers are designed for a restriction fragment of interest. All 3C primers were designed by “Primer 3”. Primers used are listed in (**Table S5**). Interactions were measured using a 3C-qPCR assay for ligation products between each anchor HindIII fragment and each target HindIII fragment. Results are presented as relative interaction frequencies compared with those GAPDH as an internal control^**28**,**63**,**64**^. After PCR, the frequency of the ligation events was estimated through agarose gel electrophoresis.

### *In vivo* Xenograft mouse model

Female 6-week-old nude mice were used. All protocols were IACUC approved, and all the mice experiments were conducted at Rincon Biosciences (Utah). Subcutaneous implantation of 17β-Estrogen pellets (0.72 mg, 60-day release; Innovative Research of America) was performed on the same day as cell injection in 6-week-old female immune-deficient nude mice. The mice were inoculated subcutaneously with 1 × 10^6^ cells of either MCF7 or MCF7TR cells, suspended in equal volumes of PBS and Matrigel. Once tumors became palpable, their sizes were measured every three days, and the tumor volume was calculated using the following formula: tumor volume (mm^3^) = length × width^2^ × 0.5. Upon reaching a mean size of approximately 150-250 mm^3^, the MCF7 and MCF7TR xenograft mice were randomly allocated into two groups, each consisting of six mice: (1) control group (treated with vehicle corn oil), (2) Brinzolamide (Selleckchem) group (oral administration at 50 mg/kg, three times a week). The data collected from this study encompassed animal weights, observations, and tumor dimensions. The data were utilized to assess drug tolerability, weight variations and gross physiological changes, as well as to evaluate anticancer activity through tumor growth inhibition or regression. Tumor volumes were monitored until the mice were euthanized. The mice were euthanized humanely, and their tumors were subsequently collected and stored for further studies.

### Differential compartments analysis

We used HiC-Pro^**22**^ to process the raw Hi-C data to generate the raw contact matrices with 100kb resolution for all tissue samples. The raw contact matrices were then fed into dcHiC^**23**^ to detect the compartment A/B and the differential compartments. Briefly, raw principal component (PC) files for each chromosome were first created by dcHiC and the best PCs within PC1 and PC2 were selected by comparing the correlation of each one against GC content and gene density. Then, the selected PCs were quantile normalized by calculating the Mahalanobis distance. Finally, four differential compartments were detected based on the normalized PCs, including two compartment transitions A-B and B-A from NTs to TTs and PTs to RTs; and the overall correlations of compartments among NTs, PTs and RTs were computed based on calculated quantile normalized PCs with a customized python script. IGV was used to visualize the compartment.

### Tumor-specific TADs analysis

We utilized HiC-Pro to process the raw Hi-C data to generate the iced contact matrices with different resolutions, including 20k, 40k, 100k and 150k, for all tissue samples. We then applied TopDom^**24**^ to call TADs for all resolutions with a series of window sizes (only parameter for TopDom), 5, 8, 10, 13 and 15. We selected the best parameters win 5, resolution 40k for TopDom based on the number of TADs and the size of TADs reported in the previous literatures^**24**,**65**^ (**Figure S2**). Since current existed tools^**66-70**^ for identifying differential TADs have several drawbacks, including only allowing a comparison of two samples, providing no specific types of TAD changes, and missing the comparison between TAD to Gap or Boundary, we developed a novel algorithm, <u>G</u>roup-, Individual-sample <u>S</u>pecific <u>TA</u>Ds (GISTA) to detect TAD changes across multiple samples. GISTA is composed of three steps: 1). Partitioning the whole genome into a series of TADs arrays across all samples; 2). Constructing a feature vector for each TADs array; 3). Annotating the type of TAD change to each TADs arrays and statistical inference. The detailed method is the following:

We denoted the sample m in n^th^ group as *G*_*k*_.*I*_*m*_ *k* ∈ and.*m* ∈ 1,2…,*M*.

1. Partitioning the whole genome into a series of TADs arrays across all samples: We applied a dynamic programming approach to identify the ‘preserved’ borders across all samples, allowing a small pre-threshold shift (default 2 bins) based on the TAD’s resolution. Then, those borders are used to partition the whole genome into a series of TADs array, noted by *TA*_*i*_, *i* = 1,2,…,*I*; *I* = *the number of broders* + 1.
2. Constructing a feature vector for each TADs array: We constructed two feature matrices for *TA*_*i*_: Neo-Del score matrix (denoted as *ND*_*i*_) and Split-Fuse score matrix (denoted as *SF*_*i*_). We defined and *nd*_*r,c*_ and *sf*_*r,c*_ (*r,c* ∈ {*G*_1_.*I*_1_,…, *G*_*k*_.*I*_*m*_})as the notes for the elements in *ND*_*i*_ and *SF*_*i*_ Before calculating *nd*_*r,c*_ and *sf*_*r,c*_, we first built a length dictionary *L*_*i*_, and a sign dictionary *S*_*i*_. The keys in *L*_*i*_ and *S*_*i*_ are samples (denoted as *s* ∈ {*G*_1_.*I*_1_,…, *G*_*k*_.*I*_*m*_}); and values are denoted as 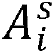 and 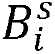, which are the length vector and the sign vector of *TA*_*i*_ for *s*, respectively. Each array element *a* in 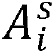 is the binsize-scaled TAD’s length, calculated by

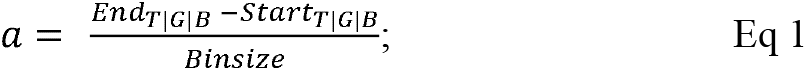

And each array element *b* in 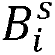 is measured by

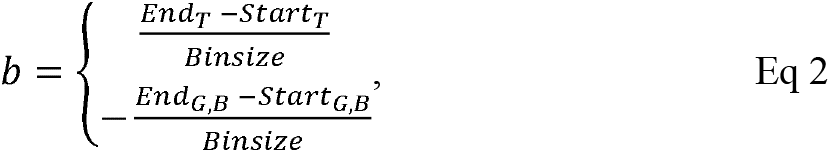

where T, G, B represent TAD, Gap and Boundary respectively, binsize is the resolution of Hi-C that used to identify TADs. Then, for element *nd*_*r,c*_ in *ND* matrix, it is then calculated by

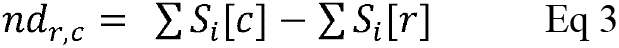

For element *sf*_*r,c*_ in *SF* matrix, it is then calculated by

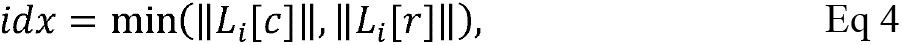

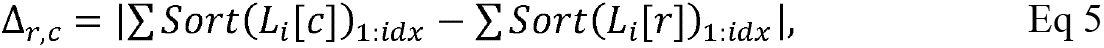

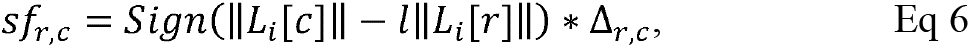

where *Sort* represents descending sort function and *Sign* represents sign function. We finally Converted *ND*_*i*_ and *SF*_*i*_ to a feature vector (denoted as *V*_*i*_) based on arithmetic mean of the pre-setting groups. For example, in this study, we have three groups, NTs, PTs, and RTs with two, five, five tumor samples, *G*_*NT*_ ⊆ (*G*_*NT*_.*I*_1_,*G*_*NT*_.*I*_2_), *G*_*PT*_ ⊆ (*G*_*PT*_.*I*_1_…*G*_*PT*_.*I*_5_), *G*_*RT*_ ⊆ (*G*_*RT*_.*I*_1_…*G*_*RT*_.*I*_5_) and we are interested in finding the *G*_*TT*_-specific TADs (*G*_*PT,RT*_ *vs G*_*PT*_), then feature vector in this case contains the following elements: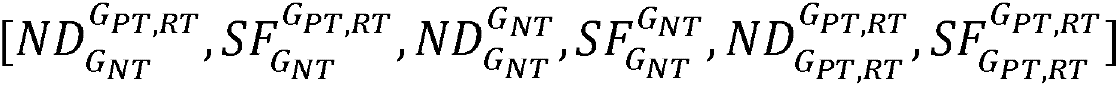, where the subscript indicates the selected columns and the superscript indicates the select rows in the *ND* or *SF*. For example, 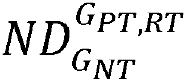 is the arithmetic mean of *ND*_*rows,cols*_, *rows* ∈ (*G*_*PT*_.*I*_1_…*G*_*RT*_.*I*_5_),*cols* ∈ (*G*_*NT*_.*I*_1_,*G*_*NT*_.*I*_2_)
3. Annotating the type of TAD changes and statistical inference: We first row-wise concatenated. all *V*_*i*_ to build a feature matrix, *F*, For the previous example (*G*_*PT,RT*_ *vs G*_*PT*_), the rows in *F* represent TADs arrays and columns are 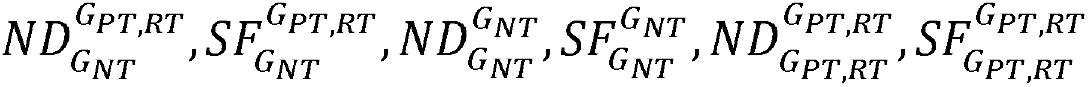. We then converted *F* to *F*_*LMH*_ by categorizng the elements in F into ‘L(ow)’, ‘M(edium)’ and ‘H(igh)’ categories. Detailly, we set *Q*_*ND*_ (0.1) and *Q*_*ND*_ (0.7) of the *F*_*I,ND*_, as well as *Q*_*SF*_ (0.1) and *Q*_*SF*_ (0.7)of the *F*_*I,SF*_ as the cutoff for ‘L’ and ‘H’, and elements besides ‘L’ and ‘H’ are ‘M’ (**Figure S3C**). In the previous example, and *F*_*I,ND*_ and *F*_*I,SF*_ are extract from *F* by selecting *ND* and *SF* related columns separately:

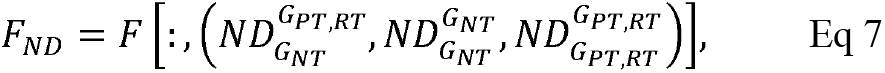

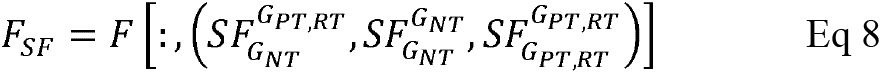

We then applied decision trees (**Figure S3D**) to annotate each TADs array to obtain distinct types of TAD changes including ‘Inter-tumor conserved TADs’ (C-TADs), ‘inter-tumor moderately variable TADs’ (MV-TADs) and ‘inter-tumor significantly variable TADs’ (SV-TADs) based on *F*_*LMH*_. C-TADs refers to TADs that are consistent across two groups, MV-TADs refers to TADs that exhibit mild variations between two groups, and SV-TADs refers to TADs with significant variations between two groups. (**Figure S3F**). We further defined six subtypes of changing within SV-TADs, which composed of the combinatorial of two individual-sample level changes, ‘individual high-specific’ (HS) and ‘individual low-specific’ (LS), as well as three TAD type changes, N(eo)D(eletion), S(plit)F(usion) and Mixed (NDSF) from *F*_*LMH*_. Specifically, HS are *TA*_*i*_ that categorized as high in either 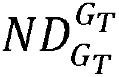 or 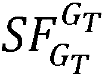,*G*_*T*_ represent Test groups and LS the complementary set of HS. ND are *TA*_*i*_ only categorized as high in 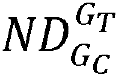, SF are *TA*_*i*_ only categorized as high in 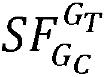 and Mixed are the rest TAs, *G*_*c*_ represents Control groups.

Finally, we performed a permutation test to infer the statistical significance of Group-Variation-H based on *F*, we first built the observations with

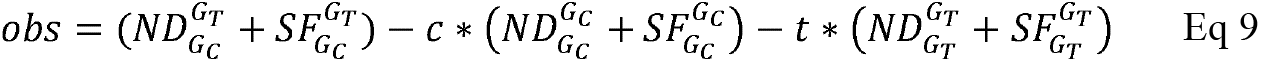

Where *c* and *t* is calculated from *argmax* (‖ *obs* < 0.05 ‖), *c, t* ∈ {0.1*n*, 0 ≤ *n* ≤ 10,*n* ∈ ℤ}. Then, the observations were permutated 1000 times. And the p values were represented by the proportion of the permutated observation that are more extreme than actual observations. The visualization of N/D/S/F scaled frequency and TADs was implemented by Logomaker^**71**^and GENOVA^**72**^, respectively.

### Looping genes analysis

The interacting loci were identified by FitHiC2^**26**^ with the parameters -r 20000 -L 40000 -x intraOnly -U 1000000. We filtered out the low interacting loci with contact counts under the 0.98 quantile of all interacting loci’s contact counts. The remaining interacting loci with high contact counts were further screened out with their genomic locations. Specifically, we defined a P-D loop as one end of an interacting loci on the promoter region of a RefSeq protein-coding gene and the other end on the distal region of the same gene. The promoter region of a gene was defined as from 4 kilobases (kb) upstream of 5’TSS to 2 kb downstream of 5’TSS. The distal region of a gene was defined as 200 kb to 10 kb upstream and downstream of 5’TSS. We denoted a loop as 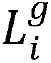, *i* ∈ {1,2,…,*N*}where N is the total number of P-D loops in a gene g. We further defined the loop intensity (*LI*) of a gene as 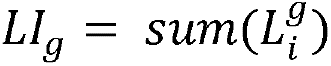. For the gene with at least one loop, we considered it as a looping gene (denoted as *LG*)

There are four stepwise process to identify four levels of looping genes. The first level is to identify individual TT-specific differential looping genes, *IDLG*_*TTs*.*vs*.*NTs*_, identified by comparing each individual TT to NTs. We denoted gained looing genes as *GLG* and lost looping genes as *LLG*. The second level is to identify individual RT-specific differential looping genes, *IDLG*_*RTs*.*vs*.*PTs*._ The third level is to identify common TR cell-specific differential looping genes with individual RT-specific differential looping genes, 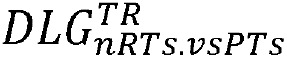, defined as the common looping genes between *IDLG*_*RTs*.*vs*.*PTs*_ and TR-specific differential looping genes. The fourth level is to identify differentially expressed looping genes, 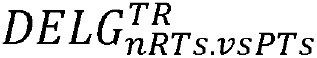, obtained by intersecting TR-specific DEGs with 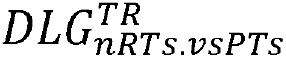. The detailed analyses were described as the following:

1. Normalizing loop intensity (*LI*) with a median of ratio method to remove the Hi-C library tissue size bias. We first constructed a loop intensity matrix (*M*_*LI*_), in which each column represents a tissue sample and each row represents a looping gene. The elements in *M*_*LI*_ were the 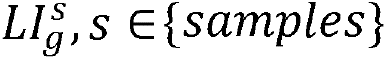 and we applied geometric average function to each row of *M*_*LI*_. All looping genes with an infinity geometric average value were filtered out and the resulting matrix was denoted as 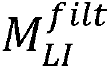. We further subtracted geometric average values of looping genes from 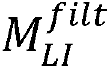 to form 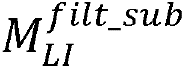 and calculated the column-based medians of 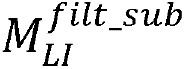. Next, we made an exponential of the medians as the normalizing factors *ceof*_*n*_ for each tissue sample. Lastly, we divided the original *M*_*LI*_ by the *ceof*_*n*_, and got the final normalized matrix 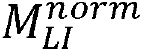.
2. Identifying *IDLG*_*TTs*.*vs*.*NTs*._ We first applied a Z-score function to each row of 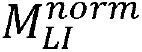; Then, we calculated signed Euclidean distance of looping genes (denoted as 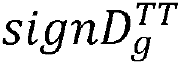) between individual tumor tissue and normal tissues:

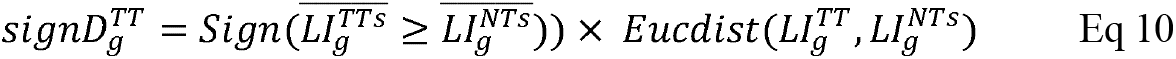

*where Sign* is a sign function, *TT* ∈*TTs* = {*PT*1,…,*PT*5,*RT*1,…*RT*5} and *NTs* = {*NT1,NT2*}. Next, we fitted the probability density function (PDF) of 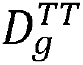 with the bi-gaussian model:

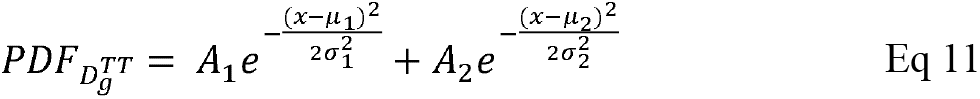

w*here* (*A*_1_,*A*_2_, μ_1_, μ_2_, σ_1_, σ_2_) are gaussian parameters that learned from the 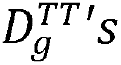 distribution; Finally, we calculated *D* value at half maximum height (HMH) of PDFs as the cutoff and identified the individual TT-specific looping genes by 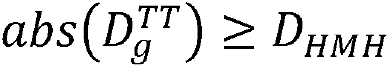
3. Identifying *IDLG*_*RTs*.*vs*.*PTs*_. We first retrieved looping genes occurred in at least two PTs (*CIDLG*_*PTs*.*vs*.*NTs*_). Then, we built a binary matrix by outer-joining *CIDLG*_*PTs*.*vs*.*NTs*_ and *IDLG*_*RTs*.*vs*.*PTs*_ as rows and {*RT*1,…*RT*5,*PT*_*c*_}as columns. 0 and 1 in the matrix represent the looping genes absent and existence, respectively. Finally, we subtracted *PT*_*c*_ column from each RT column and thus identified *IDLG*_*RTs*.*vs*.*PTs*_ when the subtracted result equal to 1.
4. Identifying 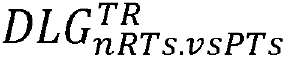. We first identified TR-specific differential looping genes, *DLG*^*TR*^, by comparing TR and TTS cell lines with the method described above. We then defined several classes of common individual RT-specific differential looping genes, denote as *DLG*_*nRT*.*vs PTs*_: *DLG*_1*RTs*.*vs PTs*_(genes shared in at least one RTs), *DLG*_2*RTs*.*vs PTs*_ (genes shared in at least two RTs), up to *DLG*_5*RTs*.*vs PTs*_ (genes shared in all five RTs). Lastly, we identified 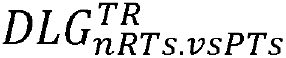 as the overlapped DLGs between *DLG*^*TR*^ and *DLG*_*nRT*.*vs PTs*_.
5. Identifying 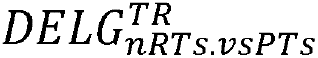. We processed DEGs in TR cell lines denoted as *DEG*^*TR*^, as described in Li .et al^**43**^ and we intersected 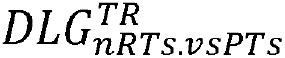 and *DEG*^*TR*^ to obtain 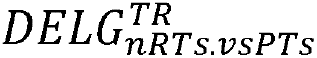. The visualization of looping genes was performed by pygenomtracks^**73**^.

### CNV analysis

We applied HiNT-CNV, a module within HiNT^**27**^, to identify CNVs from the contact matrix generated by HiC-Pro. The algorithm of CNV detection in HiNT tool utilizes a regression-based method to remove known biases and a minimizing Bayesian information criterion (BIC) procedure to get the final CNV profile. HiNT-CNV is able to take the contact matrix as the input as well as take FASTQ/BAM as the input. It outputs statistically significant CNVs. We used the default parameters in our analysis.

### Gene ontologies (GO), pathways and survival analyses

We performed the GO, pathways and survival analyses as previous described^**43**^. We applied DAVID^**29**,**74**^ and Enrichr^**30**^ to perform the pathway analysis on *LG*_*RTs*.*vs*.*PTs*_. We also used an online survival tool (www.kmplot.com)^**31**^ to analyze the prognostic value of genes on breast cancer prognosis using microarray data of 1,809 patients in terms of relapse free survival, overall survival, and distant metastasis free survival. A survival curve was displayed with the hazard ratio with 95% confidence intervals and a logrank P value.

## QUANTIFICATION AND STATISTICAL ANALYSIS

No statistical methods were used to predetermine sample sizes, but the sample sizes here are similar to those reported in previous publications. No randomization was used during data collection as there was a single experimental condition for all acquired data. Data collection and analyses were not performed blind to the conditions of the experiments as all experiments followed the same experimental condition. Statistical details of experiments and analyses can be found in the figure legends and main text above. All statistical tests were two-sided, and statistical significance was considered when P value < 0.05. To account for multiple testing, the P values were adjusted using the bonferroni correction.

