## Supplementary material for "Hi-C profiling in tissues reveals 3D chromatin-regulated breast tumor heterogeneity and tumor-specific looping-mediated biological pathways": Suppl. Materials

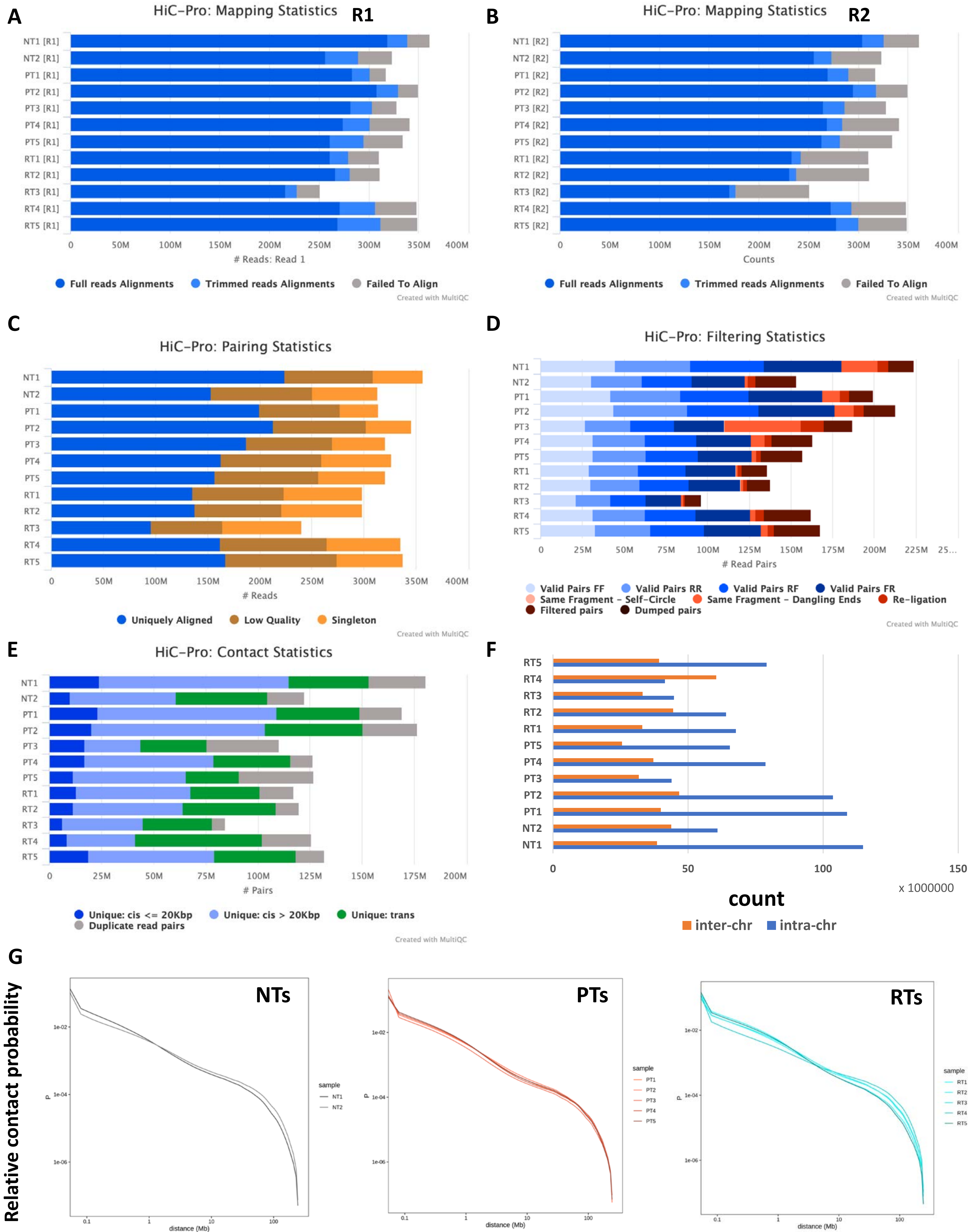

A
HiC-Pro: Mapping Statistics R1
NT1 [R1] NT2 [R1] PT1 [R1] PT2 [R1] PT3 [R1] PT4 [R1] PT5 [R1] RT1 [R1] RT2 [R1] RT3 [R1] RT4 [R1] RT5 [R1]
0 50M 100M 150M 200M 250M 300M 350M 400M
### Reads: Read 1
Full reads Alignments
Trimmed reads Alignments
Failed To Align
Created with MultiQC
B
HiC-Pro: Mapping Statistics R2
NT1 [R2] NT2 [R2] PT1 [R2] PT2 [R2] PT3 [R2] PT4 [R2] PT5 [R2] RT1 [R2] RT2 [R2] RT3 [R2] RT4 [R2] RT5 [R2]
0 50M 100M 150M 200M 250M 300M 350M 400M
Counts
Full reads Alignments
Trimmed reads Alignments
Failed To Align
Created with MultiQC
C
HiC-Pro: Pairing Statistics
NT1 NT2 PT1 PT2 PT3 PT4 PT5 RT1 RT2 RT3 RT4 RT5
0 50M 100M 150M 200M 250M 300M 350M 400M
### Reads
Uniquely Aligned
Low Quality
Singleton
Created with MultiQC
D
HiC-Pro: Filtering Statistics
NT1 NT2 PT1 PT2 PT3 PT4 PT5 RT1 RT2 RT3 RT4 RT5
0 25M 50M 75M 100M 125M 150M 175M 200M 225M 25...
### Read Pairs
Valid Pairs FF
Valid Pairs RR
Valid Pairs RF
Valid Pairs FR
Same Fragment - Self-Circle
Same Fragment - Dangling Ends
Re-ligation
Filtered pairs
Dumped pairs
Created with MultiQC
E
HiC-Pro: Contact Statistics
NT1 NT2 PT1 PT2 PT3 PT4 PT5 RT1 RT2 RT3 RT4 RT5
0 25M 50M 75M 100M 125M 150M 175M 200M
### Pairs
Unique: cis <= 20Kbp
Unique: cis > 20Kbp
Unique: trans
Duplicate read pairs
Created with MultiQC
F
RT5 RT4 RT3 RT2 RT1 PT5 PT4 PT3 PT2 PT1 NT2 NT1
0 50 100 150
x 1000000
count
inter-chr
intra-chr
G
Relative contact probability
NTs
PTs
RTs
P
1e-02 1e-04 1e-06
0.1 1 10 100
distance (Mb)
sample
NT1 NT2
PT1 PT2 PT3 PT4 PT5
RT1 RT2 RT3 RT4 RT5

### **Fig. S1**. Quality control of Hi-C data.

**A**. A stacked bar plot showing the summary of mapping status of R1 from HiC-Pro for each tissue.

**B**. A stacked bar plot showing the summary of the mapping status of R2 from HiC-Pro for each tissue.

**C**. A stacked bar plot showing the summary of the mapping pairs from HiC-Pro for each tissue.

**D**. A stacked bar plot showing the summary of the different categories of pairs from HiC-Pro for each tissue.

**E**. A stacked bar plot showing the summary of the distance of the contact from HiC-Pro for each tissue.

**F**. A bar plot showing the number of inter- and intra-chromosomal contacts for each tissue.

**G**. The RCP (relative contact probability) plots showing the classic decay line for all tissues.

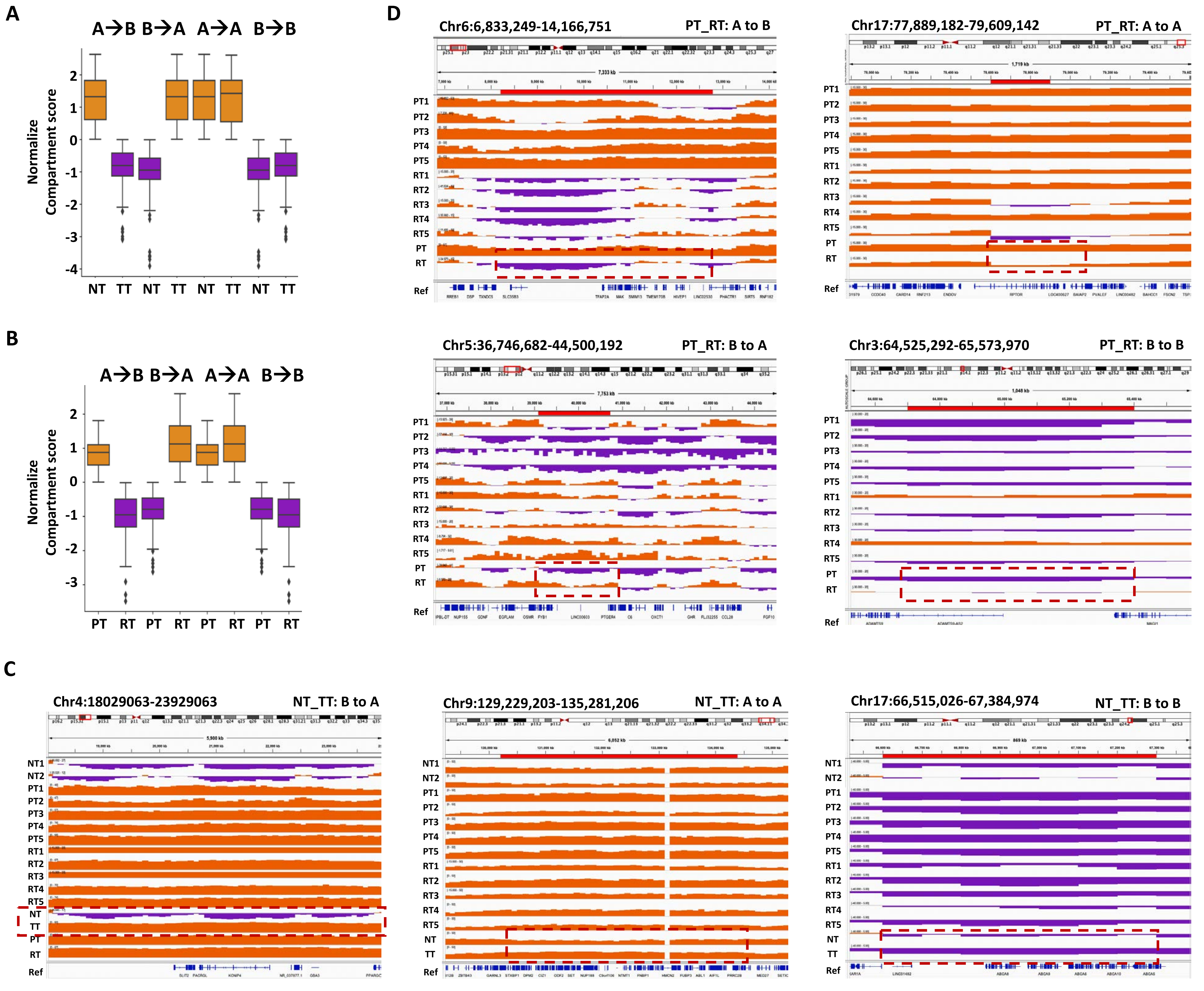
A
A→B B→A A→A B→B
Normalize Compartment score
NT TT NT TT NT TT NT TT
B
A→B B→A A→A B→B
Normalize Compartment score
PT RT PT RT PT RT PT RT
C
Chr4:18029063-23929063
NT_TT: B to A
Chr9:129,229,203-135,281,206
NT_TT: A to A
Chr17:66,515,026-67,384,974
NT_TT: B to B
D
Chr6:6,833,249-14,166,751
PT_RT: A to B
Chr17:77,889,182-79,609,142
PT_RT: A to A
Chr5:36,746,682-44,500,192
PT_RT: B to A
Chr3:64,525,292-65,573,970
PT_RT: B to B
NT1
NT2
PT1
PT2
PT3
PT4
PT5
RT1
RT2
RT3
RT4
RT5
NT
TT
PT
RT
Ref

**Fig. S2**. The identification of differential compartments among breast tumors.

**A**. The box plot showing the distribution of the normalized compartment score of the differential compartmental bins for TTs. *vs* NTs.

**B**. The box plot depicting the distribution of the normalized compartment score of the differential compartmental bins for RTs *vs* PTs.

**C**. The IGV screenshots visualizing B to A, A to A and B to B transitions for TTs *vs* NTs.

**D**. The IGV screenshots visualizing A to B, B to A, A to A and B to B transitions for RTs *vs* PTs.

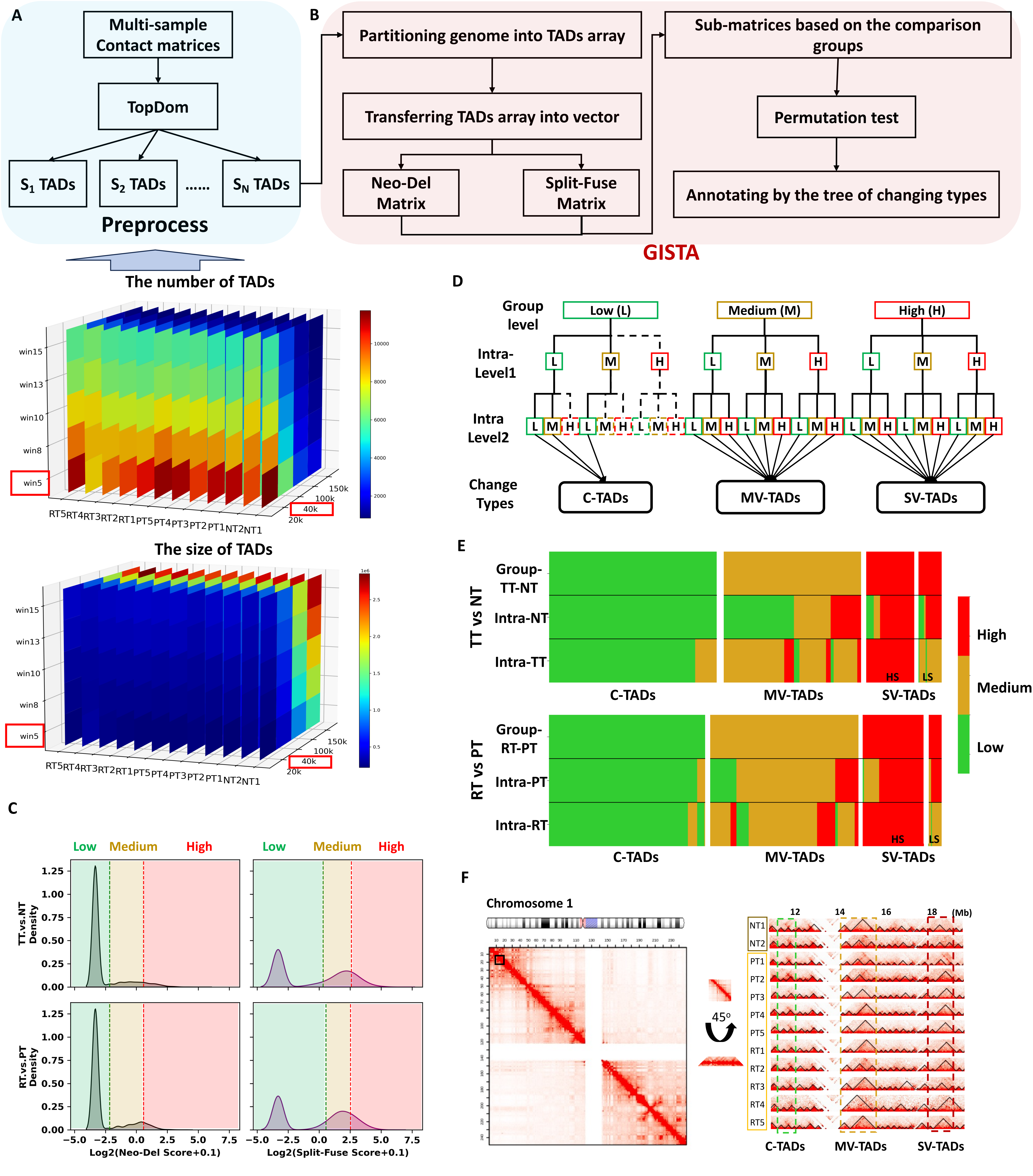

A
Multi-sample Contact matrices
TopDom
S1 TADs
S2 TADs
......
SN TADs
Preprocess
B
Partitioning genome into TADs array
Transferring TADs array into vector
Neo-Del Matrix
Split-Fuse Matrix
Sub-matrices based on the comparison groups
Permutation test
Annotating by the tree of changing types
GISTA
The number of TADs
win15
win13
win10
win8
win5
RT5 RT4 RT3 RT2 RT1 PT5 PT4 PT3 PT2 PT1 NT2 NT1
150k
100k
40k
20k
The size of TADs
C
Low
Medium
High
TT.vs.NT Density
RT.vs.PT Density
Log2(Neo-Del Score+0.1)
Log2(Split-Fuse Score+0.1)
D
Group level
Low (L)
Medium (M)
High (H)
Intra-Level1
Intra Level2
Change Types
C-TADs
MV-TADs
SV-TADs
E
TT vs NT
Group-TT-NT
Intra-NT
Intra-TT
RT vs PT
Group-RT-PT
Intra-PT
Intra-RT
HS
LS
High
Medium
Low
F
Chromosome 1
45°
12
14
16
18
(Mb)
NT1
NT2
PT1
PT2
PT3
PT4
PT5
RT1
RT2
RT3
RT4
RT5

### **Fig. S3**. The details of the algorithm, GISTA.

**A**. The workflow for group- and individual-specific TADs analysis (GISTA). Blue pad suggests the pre-processing steps by TopDom and red pad indicates the steps in GISTA.
**B**. The stacked heatmaps showing the number of TADs and the size of TADs changed with the change of two different parameters, winsize and resolution. We selected the win=5 and res=40k as the optimal parameter.
**C**. The kernel-density plot illustrating the distribution of the Neo-Del (ND) and Split-Fuse (SF) score of the TADs array. ND and SF scores were classified into three categories for the further tree annotations: low/medium/high with their 0.1 and 0.7 quantile values, represented by green, brown and red color.
**D**. A decision tree defining the types of changes for the TADs array. Three levels contain the group-level (comparisons between group1 and group2), intra-level1 (comparisons within group1) and intra-level2 (comparisons within group2). Three types of changes are inter-tumor conserved TADs (C-TADs), inter-tumor moderately variable TADs (MV-TADs) and inter-tumor significantly variable TADs (SV-TADs).
**E**. Heatmaps (Top: TT vs NT; Bottom: RT vs PT) visualizing three levels comparison for three types of changes. Each column represents a TAD array. Inter-tumor significantly variable TADs changes were further divided into individual high-specific (HS), and Individual low-specific (LS) TAD changes based on Intra-level2 values.
**F**. Left: The Hi-C contact map displaying pairwise contact frequency between genomic region across chromosome 1. Rotation of a zoom-in view by 45 degree yields a horizontal display. Right: zoom-in views of a representative region showing the three types of change of TADs array: SV-TADs (highlighted by red dash triangle), MV-TADs (highlighted by brown dash triangle) and C-TADs (highlighted by green dash triangle) for TT vs NT.

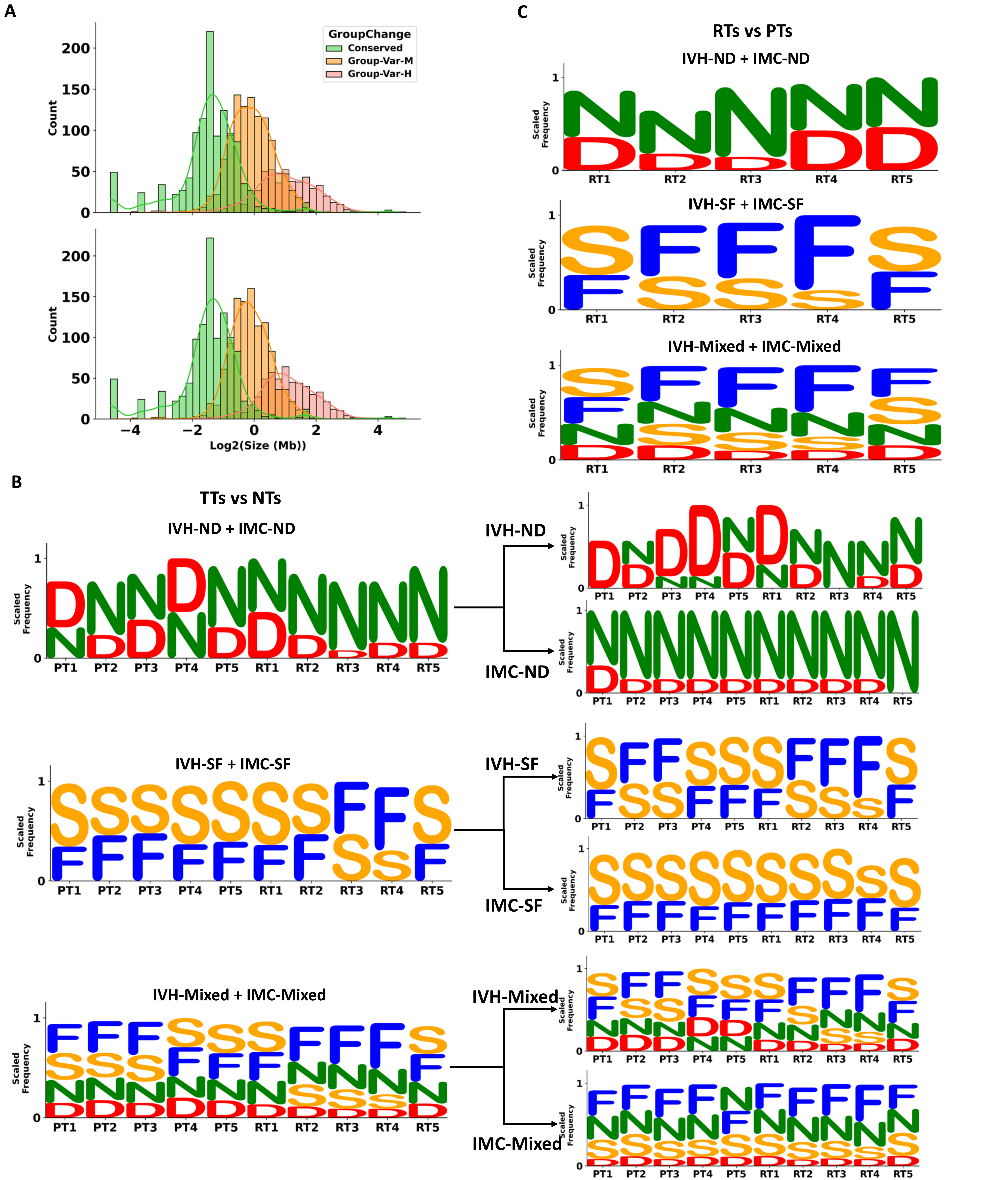
A
GroupChange
Conserved
Group-Var-M
Group-Var-H
Count
Log2(Size (Mb))
B
TTs vs NTs
IVH-ND + IMC-ND
IVH-SF + IMC-SF
IVH-Mixed + IMC-Mixed
IVH-ND
IMC-ND
IVH-SF
IMC-SF
IVH-Mixed
IMC-Mixed
Scaled Frequency
PT1 PT2 PT3 PT4 PT5 RT1 RT2 RT3 RT4 RT5
C
RTs vs PTs
IVH-ND + IMC-ND
IVH-SF + IMC-SF
IVH-Mixed + IMC-Mixed
RT1 RT2 RT3 RT4 RT5

**Fig. S4**. The frequency of four types of basic change among breast tumors.

**A**. The histograms showing the distribution of the size of three types of group-level TAD change. Top panel: TTs vs NTs; Bottom panel: RTs vs PTs.

**B**. The logo plots showing the scaled frequency of four types of basic change, N(eo), D(eletion), S(plit), F(use) for individual TTs by comparing TTs vs NTs in six sub-categories. Top: the frequency of N/D/S/F in IVH-ND and IMC-ND and their summation; Middle: the frequency of N/D/S/F in IVH-SF and IMC-SF and their summation; Bottom: the frequency of N/D/S/F in IVH-Mixed and IMC-Mixed and their summation.

**C**. The logo plots showing the scaled frequency of four types of basic change, N(eo), D(eletion), S(plit), F(use) for individual RTs by comparing RTs vs PTs in six sub-categories' summation. Top: the frequency of N/D/S/F in IVH-ND + IMC-ND; Middle: the frequency of N/D/S/F in IVH-SF + IMC-SF; Bottom: the frequency of N/D/S/F in IVH-Mixed + IMC-Mixed.

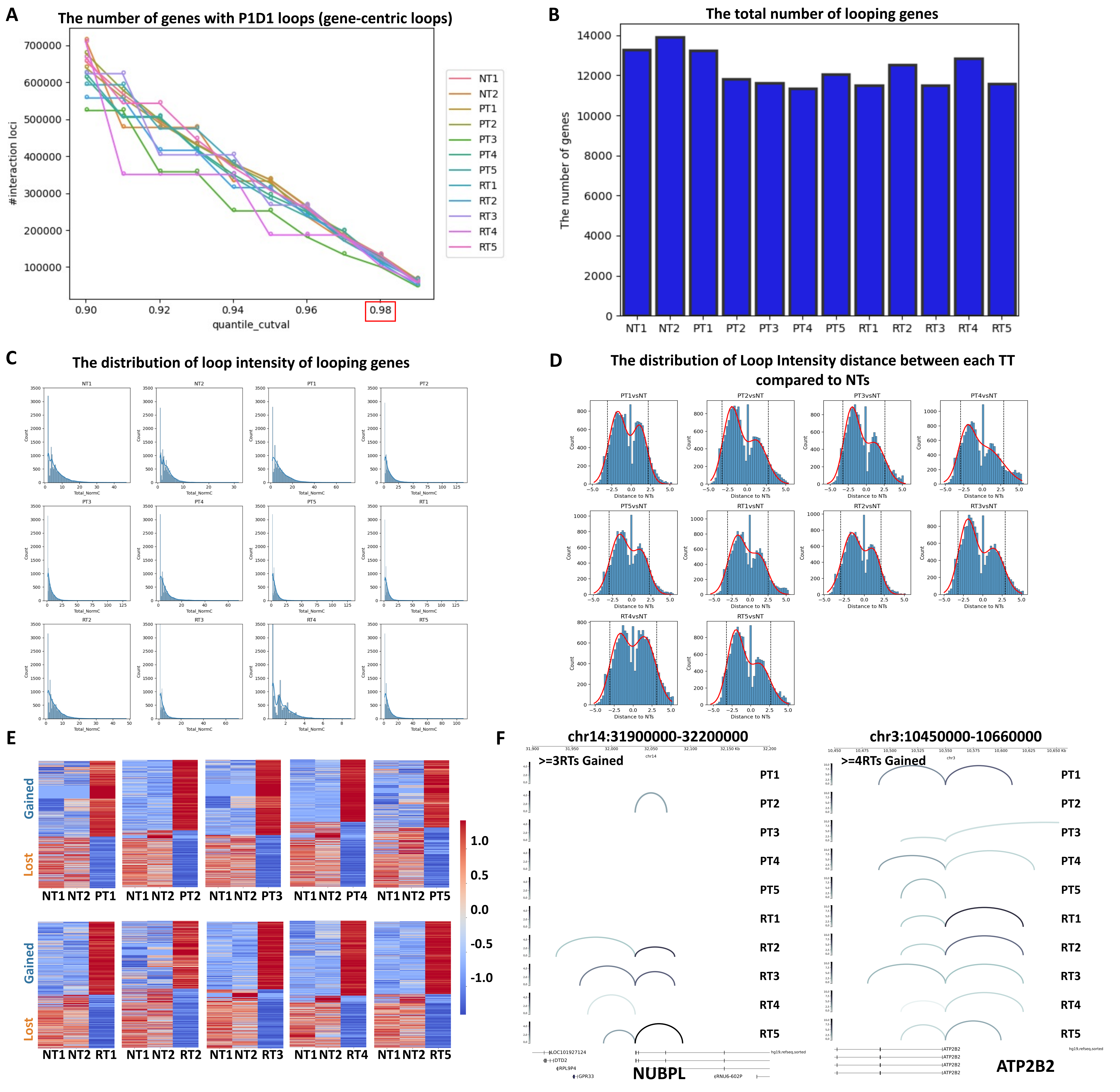

A
The number of genes with P1D1 loops (gene-centric loops)
#interaction loci
quantile_cutval
NT1
NT2
PT1
PT2
PT3
PT4
PT5
RT1
RT2
RT3
RT4
RT5
B
The total number of looping genes
The number of genes
C
The distribution of loop intensity of looping genes
Total_NormC
Count
D
The distribution of Loop Intensity distance between each TT compared to NTs
PT1vsNT
PT2vsNT
PT3vsNT
PT4vsNT
PT5vsNT
RT1vsNT
RT2vsNT
RT3vsNT
RT4vsNT
RT5vsNT
Distance to NTs
E
Gained
Lost
F
chr14:31900000-32200000
>=3RTs Gained
chr3:10450000-10660000
>=4RTs Gained
NUBPL
ATP2B2

**Fig. S5**. Identification of interaction loci, looping genes and differential looping genes among breast tumors.

**A**. The line plot showing the number of interaction loci identified at different q-value cutoffs. We used 0.98 as the final cutoff. All samples have similar level of interaction loci.

**B**. The bar plot showing the number of genes having at least a P-D loop, in which these genes were defined as looping genes (LGs).

**C**. The distribution of the looping intensity for looping genes.

**D**. The distribution of the distance of the looping intensity between TT vs NT.

**E**. The heatmaps illustrating the loop intensity of those individual tumor-specific differential looping genes.

**F**. The genome tracks visualizing the NUBPL- and ATP2B2-centric loops. The color bar ranging from 0 to 20. The darker the color, the higher the intensity of loops.

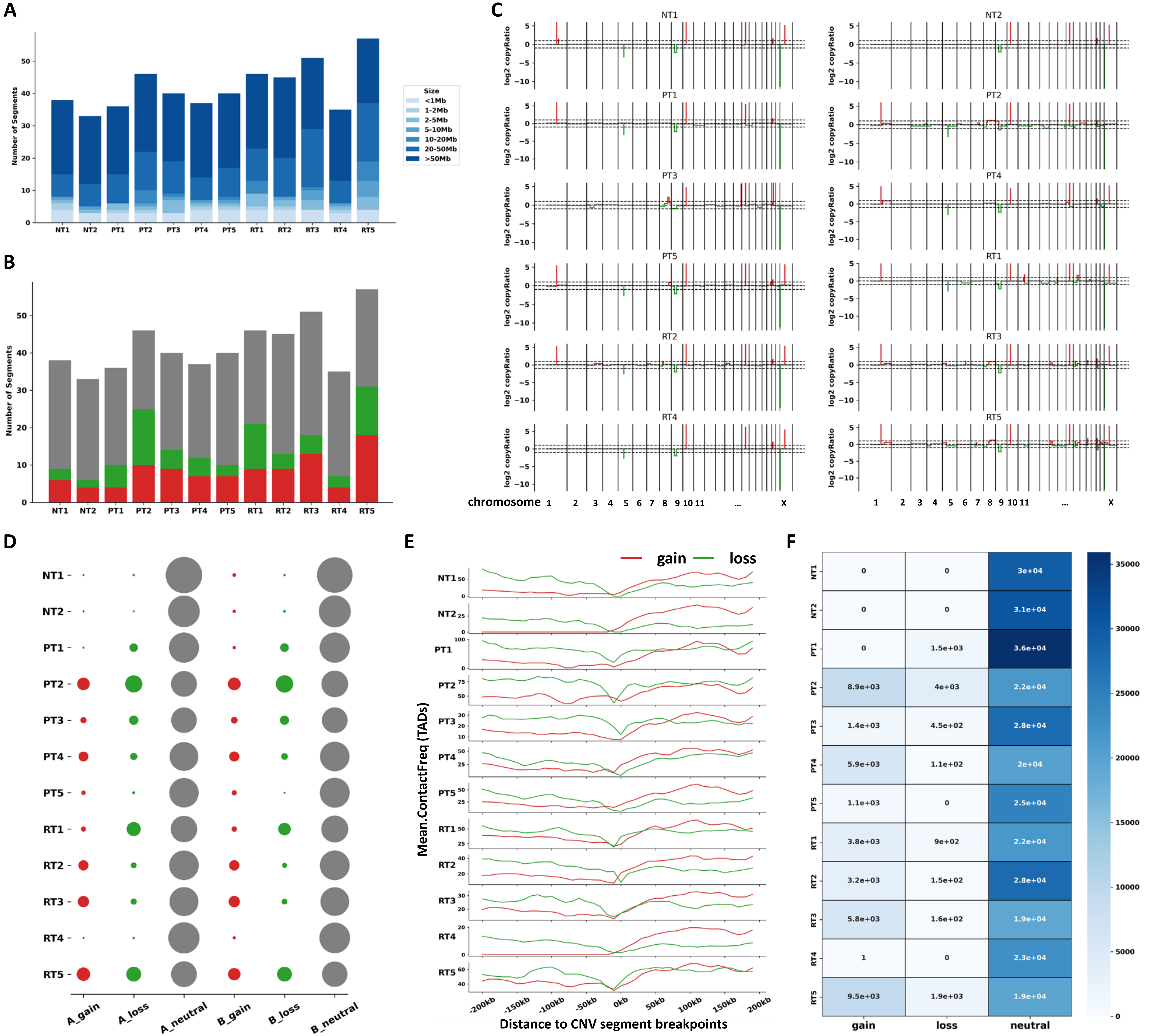
A
B
C
D
E
F
Size
<1Mb
1-2Mb
2-5Mb
5-10Mb
10-20Mb
20-50Mb
>50Mb
Number of Segments
NT1
NT2
PT1
PT2
PT3
PT4
PT5
RT1
RT2
RT3
RT4
RT5
log2 copyRatio
chromosome
1 2 3 4 5 6 7 8 9 10 11 ... X
A_gain
A_loss
A_neutral
B_gain
B_loss
B_neutral
gain
loss
Mean.ContactFreq (TADs)
Distance to CNV segment breakpoints
-200kb -150kb -100kb -50kb 0kb 50kb 100kb 150kb 200kb
neutral
NT1: 0, 0, 3e+04
NT2: 0, 0, 3.1e+04
PT1: 0, 1.5e+03, 3.6e+04
PT2: 8.9e+03, 4e+03, 2.2e+04
PT3: 1.4e+03, 4.5e+02, 2.8e+04
PT4: 5.9e+03, 1.1e+02, 2e+04
PT5: 1.1e+03, 0, 2.5e+04
RT1: 3.8e+03, 9e+02, 2.2e+04
RT2: 3.2e+03, 1.5e+02, 2.8e+04
RT3: 5.8e+03, 1.6e+02, 1.9e+04
RT4: 1, 0, 2.3e+04
RT5: 9.5e+03, 1.9e+03, 1.9e+04

**Fig. S6**. The correlation between CNVs and three layers of chromatin architecture.

**A**. The number of CNV segments (categorized by size) detected across all of the tissues.

**B**. The number of CNV segments of copy gain (log2 copy ratio > 0.3), copy loss (log2 copy ratio < -0.3) and copy neutral (log2 copy ratio between -0.3 and 0.3), respectively.

**C**. Whole genome CNV profiling across all of the tissues. Read, green and grey colors represent copy gain, copy loss and copy neutral, respectively.

**D**. The bubble plot showing the intersection results between different categories of CNV (gain, loss and neutral) and A/B compartments.

**E**. The averaged contact frequency of TADs at copy gain (red line) and copy loss (green line) CNV breakpoints.

**F**. The heatmap showing the number P-D loops overlapping with copy gain, copy loss and copy neutral CNV segments.

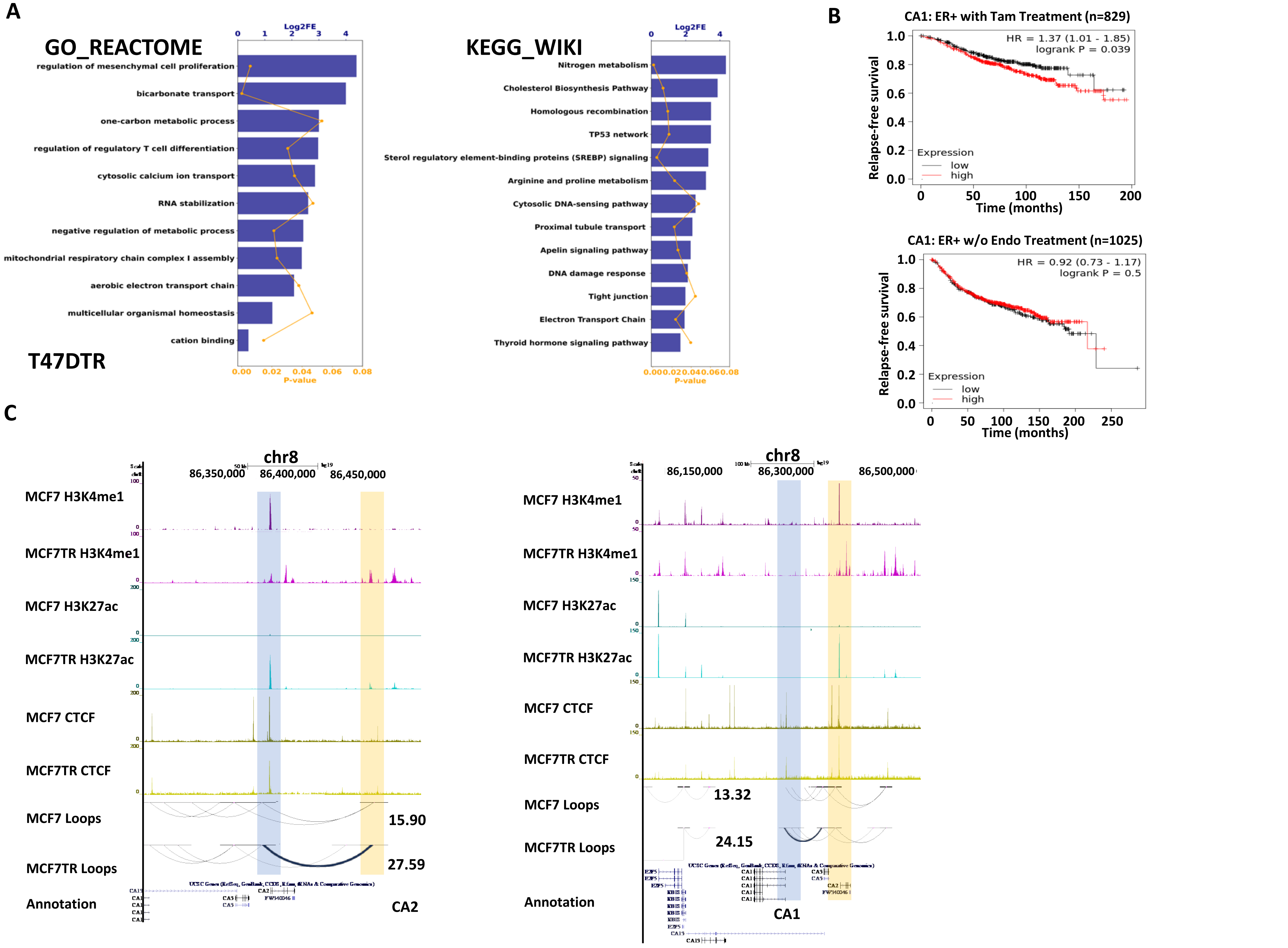
A
GO_REACTOME
Log2FE
regulation of mesenchymal cell proliferation
bicarbonate transport
one-carbon metabolic process
regulation of regulatory T cell differentiation
cytosolic calcium ion transport
RNA stabilization
negative regulation of metabolic process
mitochondrial respiratory chain complex I assembly
aerobic electron transport chain
multicellular organismal homeostasis
cation binding
P-value
T47DTR
KEGG_WIKI
Nitrogen metabolism
Cholesterol Biosynthesis Pathway
Homologous recombination
TP53 network
Sterol regulatory element-binding proteins (SREBP) signaling
Arginine and proline metabolism
Cytosolic DNA-sensing pathway
Proximal tubule transport
Apelin signaling pathway
DNA damage response
Tight junction
Electron Transport Chain
Thyroid hormone signaling pathway
B
CA1: ER+ with Tam Treatment (n=829)
HR = 1.37 (1.01 - 1.85)
logrank P = 0.039
Relapse-free survival
Time (months)
Expression
low
high
CA1: ER+ w/o Endo Treatment (n=1025)
HR = 0.92 (0.73 - 1.17)
logrank P = 0.5
C
chr8
86,350,000
86,400,000
86,450,000
MCF7 H3K4me1
MCF7TR H3K4me1
MCF7 H3K27ac
MCF7TR H3K27ac
MCF7 CTCF
MCF7TR CTCF
MCF7 Loops
MCF7TR Loops
Annotation
15.90
27.59
CA2
86,150,000
86,300,000
86,500,000
13.32
24.15
CA1

**Fig. S7**. In silico analysis of differential loping genes for T47DTR including CA1 and CA2.

**A**. Bar plots showing the results of functional analyses, including GO, REACTOME, KEGG and WIKI for >=2RTs_T47DTR differential looping genes.

**B**. Kaplan-Meier plots of CA1 showing the probability of relapse-free survival in ER+ breast cancer patients with tamoxifen treatment (n = 829) and without endocrine treatment (n = 1025). CA1 gene expression was classified as low or high (black or red lines, respectively) based on the comparison of its median cut-off value. p value was determined by the log-rank test.

**C**. The UCSC genome tracks displaying the coverage of H3K4me1, H3K27ac, CTCF, as well as the loops for CA1 and CA2 in MCF7 and MCF7TR cells.

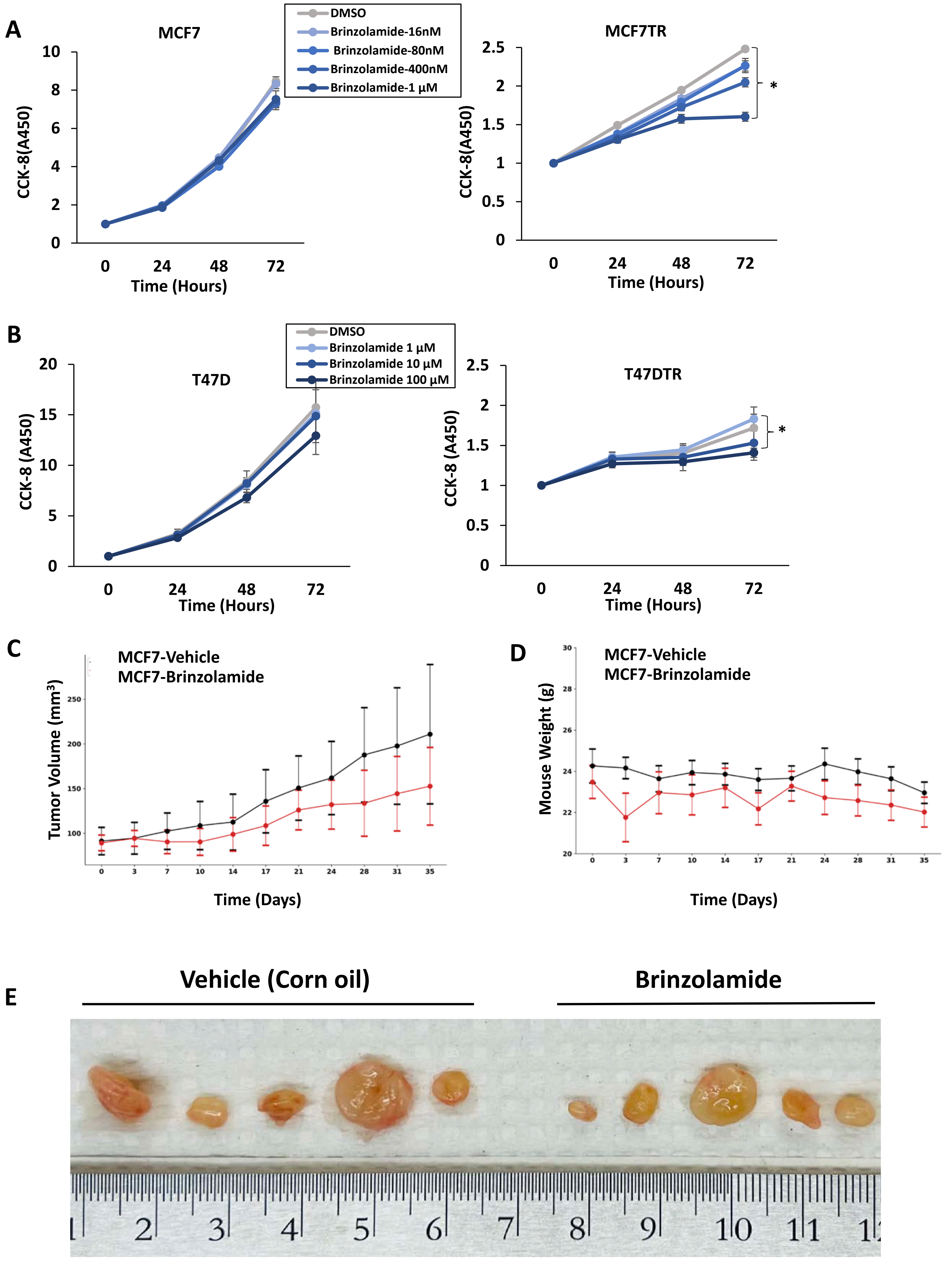
A
MCF7
DMSO
Brinzolamide-16nM
Brinzolamide-80nM
Brinzolamide-400nM
Brinzolamide-1 μM
CCK-8(A450)
Time (Hours)
MCF7TR
*
B
T47D
DMSO
Brinzolamide 1 μM
Brinzolamide 10 μM
Brinzolamide 100 μM
CCK-8 (A450)
T47DTR
C
MCF7-Vehicle
MCF7-Brinzolamide
Tumor Volume (mm3)
Time (Days)
D
Mouse Weight (g)
E
Vehicle (Corn oil)
Brinzolamide

**Fig. S8.** The effect of CA2 inhibitor, Brinzolamide in MCF7 cell growth *in vitro* and *in vivo*.

**A.** Time and dose dependent cell growth after Brinzolamide treatment in MCF7 and MCF7TR cells by CCK-8 assay. MCF7 and MCF7TR cells were seeded at 1000 cells per well in a 96-well plate and treated with different concentrations of Brinzolamide. Values are expressed as the mean ± Standard deviation of three independent experiments. Samples t-test, *p < 0.05.
**B**. Time and dose dependent cell growth after Brinzolamide treatment in T47D and T47DTR cells by CCK-8 assay. T47D and T47DTR cells were seeded at 1000 cells per well in a 96-well plate and treated with different concentrations of Brinzolamide. Values are expressed as the mean ± Standard deviation of three independent experiments. Samples t-test, *p < 0.05.
**C**. Effects of Brinzolamide on MCF7 cell-derived Xenograft mice. When tumors became palpable, tumors were treated with control, Brinzolamide. Tumor growth was analyzed by measuring the tumor volume. Error bars represent standard error mean. Significance is shown only for endpoint measurements.
**D**. Mice weights over time for each treatment group. Error bars represent standard error mean.
**E**. Effects of Brinzolamide on MCF7 cell-derived Xenograft mice. Photographs of mammary tumors for each treatment group at the study endpoint; ruler scale is mm.
